# Supra-molecular assemblies of ORAI1 at rest precede local accumulation into punctae after activation

**DOI:** 10.1101/2020.01.13.903856

**Authors:** Diana B. Peckys, Daniel Gaa, Dalia Alansary, Barbara. A. Niemeyer, Niels de Jonge

**Affiliations:** Molecular Biophysics, University of Saarland, Center for Integrative Physiology and Molecular Medicine, 66421 Homburg/Saar, Germany; INM – Leibniz Institute for New Materials, 66123 Saarbrücken, Germany; Department of Physics, University of Saarland, 66123 Saarbrücken, Germany

## Abstract

The Ca^2+^ selective channel ORAI1 and endoplasmic reticulum (ER)-resident STIM proteins form the core of the channel complex mediating store operated Ca^2+^ entry (SOCE). Using liquid phase electron microscopy (LPEM) the distribution of ORAI1 proteins was examined at rest and after SOCE-activation at nanoscale resolution. The analysis of over seven hundred thousand of ORAI1 positions showed that already at rest, a majority of the ORAI1 channels formed STIM-independent distinct supra-molecular clusters. Upon SOCE activation and in the presence of STIM proteins, ORAI1 assembled in micron-sized two-dimensional (2D) structures, such as the known punctae at the ER plasma membrane contact zones, but also in divergent structures such as strands, and ring-like shapes. Our results thus question the hypothesis that stochastically migrating single ORAI1 channels are trapped at regions containing activated STIM, and we propose instead that supra-molecular ORAI1 clusters fulfill an amplifying function for creating dense ORAI1 accumulations upon SOCE-activation.

**STATEMENT OF SIGNIFICANCE:** ORAI1 proteins form channels mediating store operated Ca^2+^ entry, an important trigger for many cellular functions, especially in the immune system. ORAI1 channels at rest are assumed to be randomly distributed in the plasma membrane, while they accumulate into so-called “punctae” upon activation, where binding by STIM proteins activate the Ca^2+^ channels. Using liquid phase electron microscopy, we discovered that ORAI1 forms small, elongated clusters indicating the existence of supramolecular assemblies. The role of such supramolecular organization of ORAI1 is possibly an amplifying function for the effective creation of ORAI1 accumulations in punctae, since the binding of only one ORAI1 protein would trap a multiple of channels.

## INTRODUCTION

The molecular processes preceding and optimizing the activation of store-operated calcium entry (SOCE) are difficult to study because they require single molecule resolution, while at the same time, the interaction of a multiple of proteins needs to be examined (1, 2). After several early reports supported tetrameric ORAI1 conformations (3-6) the current view is that ORAI1 proteins assemble as a hexameric channel complex at rest (7-9), and that these hexameric protein complexes are randomly distributed throughout the plasma membrane. Upon Ca^2+^ store depletion, STIM proteins redistribute in the endoplasmic reticulum (ER) membrane towards the cytosolic side of ER plasma membrane contact zones, where typical, dense STIM1 accumulations, so-called punctae are formed (10, 11). In these junctional areas between ER and plasma membrane, STIM1 proteins approach the plasma membrane on the cytoplasmic side where they interact with ORAI1, resulting in a conformational change that opens the ORAI1 channels. Yet, how the ORAI1 channels translocate to these regions in order to get trapped there is not fully understood. It is assumed that single ORAI1 channels randomly diffuse within the plasma membrane, until they arrive at sites of activated STIM1, so-called punctae (2). Here, ORAI1 binds to STIM1, leading to an accumulation of ORAI1 in the plasma membrane, above the STIM punctae, thus mirroring the distribution pattern of the punctae but then at the cell exterior, accompanied by activation of the Ca^2+^ channels, resulting in a local Ca^2+^ influx (2). But possibly different mechanisms play a role, for example, the additional insertion of ORAI1 from intracellular stores into the regions of punctae (12), or potentially a pre-clustering of several hexamers which are then pulled in or trapped at the ER-PM junctions.

By using liquid phase electron microscopy (LPEM) capable of studying ORAI1 proteins in the plasma membrane of intact cells (9, 13), we set out to examine the difference in spatial distribution of ORAI1 between cells at rest and upon SOCE-activation. Our aim was to test 1) if single ORAI1 channels are indeed randomly distributed at rest, and then move into clusters as independent entities, and 2) if activated clusters show distinct patterns. For this purpose, ORAI1 proteins with a stoichiometric expression of cytosolic GFP and with a human influenza (HA) tag located in the extracellular loop between transmembrane regions (TM) 3 and 4, were expressed in HEK cells, and labeled with quantum dots providing both a fluorescence label for light microscopy, and a nanoparticle label for detection with electron microscopy (9, 13). Fig. 1a depicts the dimensions of an ORAI1 hexamer (7) in a hypothetical arrangement of three QD labels bound to the same ORAI1 hexamer. Due to the flexibility of the linker, the center-to-center distance between both QDs may vary between 20 and 40 nm. Spatial patterns of individual QD labels on whole cells were imaged under resting conditions, and after two methods of SOCE-activation. The spatial distributions of the labeled ORAI1 proteins were subsequently analyzed with nanometer spatial resolution, as obtained with the LPEM method (14, 15).

**FIGURE 1.**
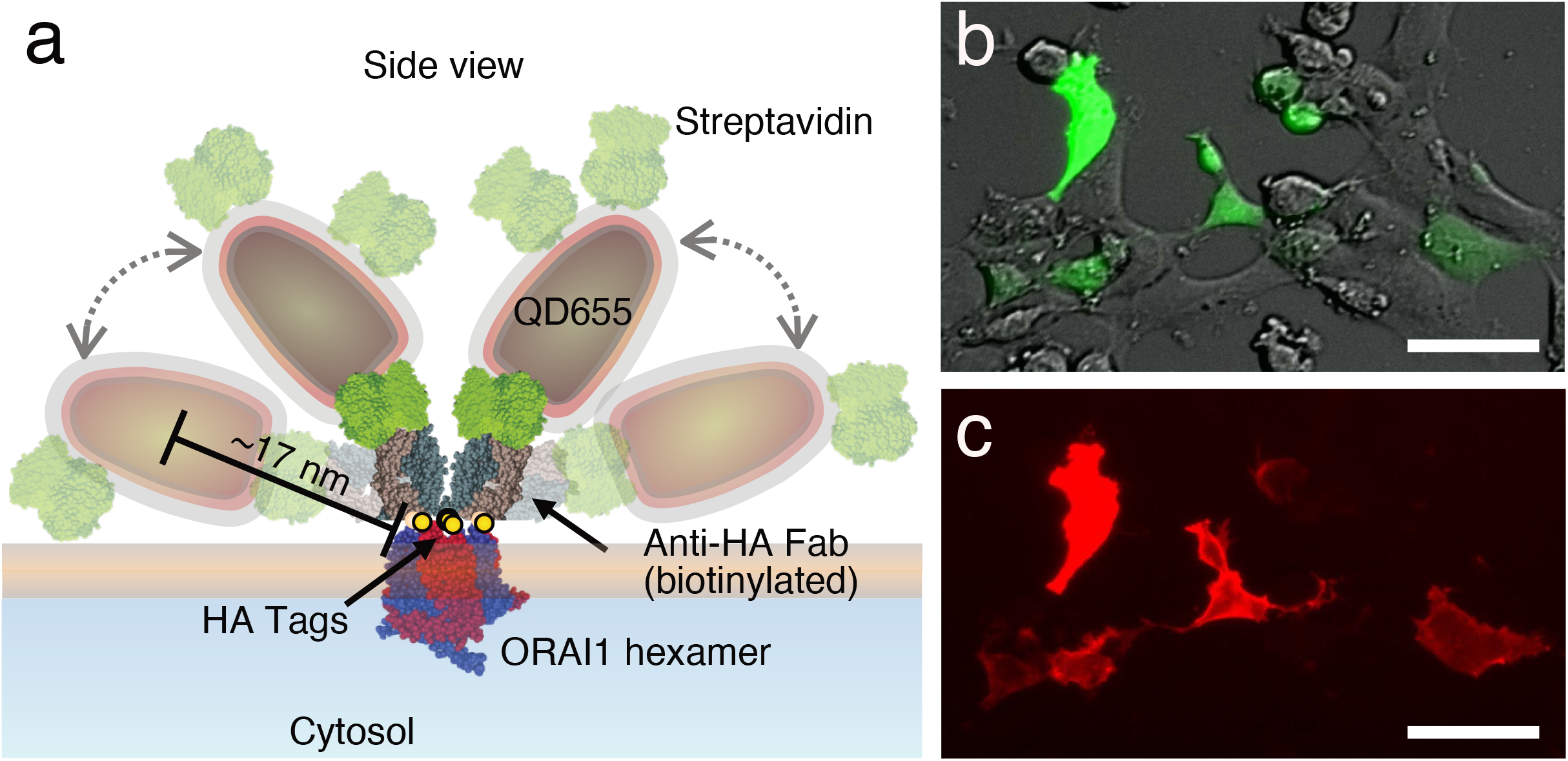
Labeling of ORAI1 proteins in the plasma membrane of HEK cells with quantum dots (QDs). **(a)** Schematic representation of QD-labeling of the ORAI1 Ca^2+^ channel in the plasma membrane. Labels consisted of a biotinylated Anti-HA Fab that bound to the HA-tags (yellow spheres) at the extracellular region of each subunit of the ORAI1-HA channel, shown in hexameric conformation (blue and red). The bound Fab was labeled using streptavidin (green) conjugated, fluorescent QDs with an electron dense core (dark gray), and an electron transparent polymer coating shell (yellow). Shown are several possible spatial positions of two exemplary bound QDs, the positions showing half-transparent labels represent a maximally extended configuration. The drawn-to-scale representation of all involved molecules, reveals a maximum distance between the center of a bound QD and the ORAI1 hexamer of 17 nm. All protein structures were derived from the Protein Data Base, sizes of QDs were measured elsewhere (9) all structures are drawn to scale. **(b)** Overlay of direct interference contrast light microscopy, and fluorescence microscopy of green fluorescent protein (GFP) intracellularly co-expressed with the same ratio as ORAI1. **(c)** Fluorescence images of QD-labeled ORAI1 at rest showing overlap with the cells expressing ORAI1 visible from GFP in (b). Scale bars = 50 μm.

## MATERIALS AND METHODS

### Materials

Fetal bovine serum, 2-mercaptoethanol (ME), thapsigargin (Tg), cyclopiazonic acid (CPA), and sodium azide, were either from Fisher Scientific or Sigma Aldrich. ScreenFect^®^A Transfection reagent was from Incella GmbH, Eggenstein-Leopoldshafen, Germany. Anti-HA-biotin, high affinity (3F10) from rat IgG1 was from Roche Diagnostics, Mannheim, Germany. Dulbecco’s phosphate buffered saline (DPBS), Modified Eagle’s Medium (MEM), normal goat serum (GS), CellStripper and quantum dot Qdot^®^ 655, and Qdot^®^565 streptavidin conjugates (QD) were from Fisher Scientific GmbH, Schwerte, Germany. ROTISOLV^®^ high pressure liquid chromatography grade pure water, acetone and ethanol, phosphate buffered saline (PBS) 10 × solution, electron microscopy grade glutaraldehyde (GA) 25% solution, D-saccharose, sodium chloride, glycine, biotin free and molecular biology grade albumin fraktion V (BSA), and sodium cacodylate trihydrate were from Carl Roth GmbH + Co. KG, Karlsruhe, Germany. Electron microscopy grade formaldehyde (FA) 16% solution was from Science Services GmbH, Munich, Germany. 0.01% poly-L-Lysine (PLL) solution (mol wt. 70,000-150,000), sodium tetraborate, and boric acid were from Sigma-Aldrich Chemie GmbH, Munich, Germany. CELLVIEW cell culture dishes (35 mm) with four compartments and glass bottoms were from Greiner Bio-One GmbH, 72636 Frickenhausen. Custom designed silicon microchips were purchased from DENSsolution, Netherlands. The microchips had outer dimensions of 2.0 × 2.6 × 0.4 mm^3^ and each contained a silicon nitride (SiN) membrane window, usually with dimensions of 150 × 400 μm^2^ (sometimes larger) along with a membrane of 50 nm thickness. Trivial transfer multilayer graphene was purchased from ACS Material LLC, Pasadena, CA, USA. NaCl2 crystals were from Plano GmbH, Wetzlar, Germany.

HEK293 cells were obtained from ATCC and genetically modified using CRISPR/Cas9-mediated gene deletion of endogenous ORAI1-2 genes (named CRI_1), or ORAI1-3 genes (named CRI_2, which were used for controls shown in the supplementary information), as described in (9). A HEK cells line lacking endogenous STIM proteins (CRI_STIM) (16) was obtained from was obtained from Donald L. Gill (16). An ORAI1 construct with an extracellular, nine amino acid HA-tag within the second extracellular loop of ORAI1, was used for labeling the protein with a QD (17). The DNA construct contained also DNA encoding green fluorescent protein (GFP) but separated by a cleavable peptide sequence (P2A) thus guaranteeing the same expression ratio (13, 18).The ORAI1-GFP DNA construct was transiently expressed in the HEK cells using ScreenFect^®^A as described in (9), which also includes the results of functional tests of the transfected ORAI1-HA DNA constructs.

### Preparation of microchips with transfected HEK cells

CELLVIEW dishes and microchips with thin SiN windows were used as a support for CRI_1 cells. Preparation of new microchips was performed as described previously (19), briefly, protective photoresist coating on the microchips was removed with acetone and ethanol, the microchips and dishes were plasma-cleaned for 5 minutes, coated with PLL, and immersed/filled with cell medium (supplemented with 10% FBS and 50 μM ME). Cells grown in a 25 cm^2^ flask were harvested at ~90% confluency, and washed once in supplemented cell medium. Cell transfection was performed according the supplier instructions. In experiments with ORAI1 at rest, 0.25 μg ORAI1-HA DNA, and 1.5 μL transfection reagent (TR) were used per compartment of a 4-compartment dish (35 mm diameter) containing a final volume of 420 μL cell suspension. In experiments with activated ORAI1, 0.17 μg ORAI1-HA DNA, 0.55 μg STIM1-mcherry DNA, to yield a 1:3 ratio between ORAI1 and STIM1, or 0.17 μg for each DNA and 2 μL TR, for a 1:1 ratio, were used per dish compartment. The respective volumes for single microchips kept in wells of a 96-well plate were 25% of those used for dish compartments. The cell samples were then incubated for 24 hours in the CO_2_ incubator.

### Labeling of overexpressed ORAI1-HA at rest or after activation

A two-step labeling protocol (13, 19) was applied using a biotinylated anti-HA Fab, followed by labeling with 20 nM streptavidin-conjugated QD, as previously described (9). After transfection, the samples were rinsed with supplemented cell medium pre-warmed to 37°C. For experiments examining unstimulated ORAI1 distribution, in the following named cells at rest, the samples were briefly rinsed in pre-warmed 0.1 M cacodyl buffer containing 0.1 M sucrose, pH 7.4 (CB), and incubated in 3% formaldehyde/0.2% glutaraldehyde in CB for 10 minutes at room temperature, thereby assuring fast and permanent fixation of membrane proteins, eliminating their diffusion (21-23). For the examination of activated ORAI, cells were first rinsed twice with pre-warmed medium (MEM, suppl. with 10% FCS and 50 μM ME), then incubated with 1 μM Tg, or 30 μM CPA (both in supplemented medium), for 15 min at 37 °C (in the CO2 incubator). The activated cells were rinsed once with pre-warmed medium, and once with pre-warmed CB, followed by fixation as described above. Fixation was terminated by rinsing once with CB, three times with PBS, 2 min of incubation in GLY-PBS (0.1% glycine in PBS) for 2 minutes, followed by a rinse in PBS. The cells were then incubated in 400 ng/mL Anti-HA-Fab-Biotin lab. sol. in PBS, first for 1 hour at room temperature, followed by 3 – 6 hours at 4°C. The QD-labeling solutions were prepared by first diluting 1 μM Streptavidin-QD stock solutions 1: 5 in 40 mM Borate buffer, pH 8.3, and a further dilution in hBSA-PBS (PBS with 1% BSA) to obtain a 20 nM QD labeling solution. After three times rinsing in PBS, cells were incubated in the streptavidin-QD labeling solutions for 12 minutes at room temperature. Since no unspecific binding occurred without addition of BSA, it was omitted in the processing steps before the QD-incubation. After the QD incubation, the cells were rinsed four times with hBSA-PBS before fluorescence microscopy was performed.

### Fluorescence Microscopy

After QD incubation and before the second fixation step, cells on microchips were imaged with an inverted fluorescence microscope (DMI6000B Leica, Germany) in a pristine, 35 mm cell culture glass-bottom dish filled with 2 mL hBSA-PBS. 20x and 40x objectives were used together with four channels, one collecting direct interference contrast (DIC), and three fluorescence channels for, GFP (460 – 520 nm excitation, 515 – 560 nm emission), QD655 (340 – 380 nm excitation, >420 nm emission), and QD565 (540 – 560 nm excitation, 580 – 620 emission).

### Processing of samples for LPEM

To stabilize the cells on the microchips for electron microscopy, the aforementioned samples were further fixed with 2% glutaraldehyde in CB for 10 minutes at room temperature. After one rinse with CB, and three rinses with hBSA-PBS, they were stored in hBSA-PBS supplemented with 0.02% sodium azide, at 4°C until liquid-phase STEM, usually performed within 1-3 weeks. To keep the ORAI channels in their almost native environment, as provided by the imaging of hydrated, intact cells, the microchip samples were covered with graphene. Multi-layer (3 to 5 layers) graphene on polymer was cleaned and transferred onto the sample as described previously (15, 24). For the coating of a microchip sample, a graphene sheet of approximately the size of a microchip was detached from its supporting NaCl crystal through immersion in a beaker filled with HPLC-grade water, placed under a binocular. The (wet) microchip was grabbed with a pair of fine-tipped tweezers, rinsed twice with pure water, and immersed in the liquid below the floating graphene. The graphene was then carefully scooped up by slowly drawing the microchip up and out of the water. The tweezers tips, still holding the microchip with the on top swimming graphene sheet, were fixed with a small rubber O-ring, and the other tweezer end was clamped into a small stand, so that the microchip hang free in the air. After a few minutes dry time the water on the graphene had evaporated and the graphene was directly adhering to the underlying, still hydrated cells, a detailed description of this step can be found elsewhere (15).

### LPEM of QD-labeled whole cells

To observe the individual QD-labeled ORAI1-HA positions, the graphene-coated samples were imaged in a transmission electron microscope (ARM 200, JEOL, Japan), with STEM dark field mode (15). The following settings were used: electron energy of 200 kV, and 175 pA probe current. For orientation purpose, the imaging session started with the recording of two to three overview STEM images, covering the entire SiN-window area. These low magnification images were directly compared to the previously recorded fluorescence- and DIC images and served to navigate to selected, representative cells chosen for high-magnification imaging. Several images from selected QD-labeled cells were recorded from randomly chosen plasma membrane regions. Note that central areas in the thickest, nucleus containing part of the cells were excluded. Images of QD655 labeled cells were recorded with magnifications of mostly 60.000×, respect. 120.000× for QD565 labeled cells (occasionally, other magnifications ranging between 40,000 and 150,000× were applied), resulting in a pixel size of 1.6 nm for QD655 images, respect. 0.8 nm for QD565 images. The image size of the 16-bit images was 2048 × 2048 pixels, comprising a scanning area of 10.1 μm^2^ per 60.000× magnified image, respect. 2.9 μm2 per 120.000× magnified image. The used pixel dwell-times ranged between 8 and 14 μs. The calculated electron doses for images of cells labeled with QD655 was usually between 40 and 60 e^-^/Å^2^, cells labeled with the smaller QD565 (shown in the supplementary information) had maximally a dose of 125 e^-^/Å^2^, which is below the reported limit of radiation damage for such samples (25).

### Particle detection

In order to obtain the lateral coordinates of the QD labels, all STEM images were first visually screened for infrequently occurring contaminants on the graphene (remnants of the production process) with dimensions and contrast characteristics sometimes hampering the automated label detection. In such cases, ImageJ (version 1.52a, NIH) was used to manually blank the respective contaminant in the image by covering them with a fitting shape, filled with the grey value of the surrounding background. QD labels were detected and localized by applying a dedicated Plugin of local design in ImageJ described elsewhere (26). The main processing steps consisted of a Fourier filter for spatial frequencies between a factor of 3 smaller and a factor of 3 higher than the set size (7 nm), and a binarization with an automated threshold with a maximum entropy setting. The particles were automatically detected using the “Find Particles” tool, with a precision corresponding to the pixel size of 1.6 nm. A demonstration of the particle detection technique can be found in an earlier publication, including also an error estimation (27).

For a better visual perception of the arrangement of labeled ORAI in selected images, we implemented another Plugin of local design for the purpose of improved visual pattern recognition of the arranged labels. This was achieved by using the x-y center position data from the detected QD labels in a STEM image, and drawing black circles around each center position, on a blank image background of the same dimensions as the STEM image. Each circle thereby indicated the plasma membrane area where the underlying ORAI1 protein and bound labels localized. The size of the circle was determined by the dimensions of the QDs and the flexible linker, including the conjugated streptavidin, and the Anti-HA Fab. As can be seen in the drawn-to-scale model shown in Figure 1 and Fig. S1, the maximal distance between the center of a QD and the ORAI1 protein is ~17 nm, so that the diameter of the circle was set to 35 nm.

### Analysis of supra-molecular ORAI1 arrangements and punctae

For the detection of possible ORAI1 supramolecular arrangements, the processed STEM images, showing the labels as 35 nm diameter black circles, were analyzed with the “Analyze Particles” tool of ImageJ. The size limit was set such that clusters consisting of >2 labels were detected. The result was visually inspected and, if necessary corrected, for a few mistakenly detected clusters with only two circles. All fully depicted clusters >2 were analyzed for their major and minor axis length, as well as for their aspect ratio, with the “Measure” tool.

For the quantitative analysis of ORAI1 distribution in punctae, all recorded images from all maximally SOCE-activated cells were visually inspected in order to identify images displaying punctae. These punctae were defined as regions of locally accumulated labeled ORAI1 with at least 2-times higher label densities than in the surrounding regions, limited by a visually identifiable border towards the surrounding regions of lower ORAI1 density. Accumulated ORAI1 areas of strand- or ring-like shapes were excluded from the analysis, due to the difficulty of defining their borders. 58% of all recorded images from the group of maximally SOCE activated cells contained such punctae. With the “Freehand selection” tool in ImageJ all fully depicted punctae were manually marked at their borders, thus yielding regions of interest (ROI). The major and minor axis length, as well as their area size of these ROI was determined with the “Measure” tool. Thereafter, the labels inside the ROIs were blanked, by filling the ROIs with grey background color, and the remaining number of labels in the regions outside the ROIs was again determined using the “Find particles” tool, similarly as done previously for all recorded images (as explained in the previous paragraph). The difference between the number of particles detected outside the ROI and the total number of particles in the respective image yielded the number of particles within the ROI.

### Simulation of label distributions corresponding to selected STEM image data sets

Spatial label distributions were simulated to discern if the detected label clusters >2 were bound to supramolecular ORAI1 arrangements, consisting of more than one underlying ORAI1 hexamer, or if similar clusters could also derive from randomly distributed hexameric ORAI1. These simulations were based on data recorded from different experimental groups, each consisting of a set of 12 images from cells displaying low ORAI1 expression levels, and a set of 6 images from cells with high expression levels. For every image a corresponding simulation was made, using the same number of detected particles and the image size, but the distribution of the labels was determined by an algorithm based on randomly scattered hexameric ORAI1, labeled with a 30% labeling efficiency, and including effects of spatial hindrance of a newly bound QD to already bound QDs using a model of hard shell QDs with a surrounding soft shell. These simulated label distributions were processed and analyzed in the same way as the data from the STEM images.

When creating a new particle simulation, as a first step, a given number of virtual ORAI1 proteins were placed randomly within the image. Their positions followed a uniform distribution while overlapping proteins were strictly avoided by limiting the minimum distance between hexamers to 10 nm, representing the largest diameter of the protein complex located intracellularly. Each of the proteins was then loaded with up to six circular labels. The exact number of labels at each protein was determined by a random process modelling the labeling efficiency. Also, the center-to-center distance between protein and label was generated randomly but restricted to a maximum of 20 nm. The directions of the displacements were generated randomly. Overlap of the hard shell of the labels (5 nm radius) was prohibited, i.e. the minimal distance between two label centers was 10 nm.

Additionally, the soft shell of the labels (additional 3,5 nm radius) was modeled by sometimes allowing and sometimes prohibiting an overlap. The probability for accepting a particle’s position when soft shells were overlapping was calculated as:

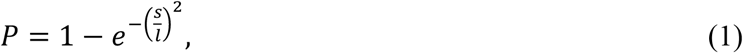

With soft shell thickness *l* = 3.5 nm, and the randomly generated distance between the hard shells *s* ∈ [0, 2×1]. This generated minimal acceptance probability if an overlap of the entire soft shell occurred, i.e. the hard shells touched (*s* = 0). For soft shells that were barely touching (*s* = 2×*l*), the probability was maximized.

If a generated label position led to an overlap (as described above) with a previously placed label, a new position was created. If necessary, this was repeated multiple times before the label was discarded entirely, assuming that no valid position would be found in this area. The maximum number of attempts was 50. To avoid any bias, the labels were placed sequentially, starting with the first label of every single protein before continuing with the next ones. Finally, a simulated image was created from these label positions using the aforementioned mentioned ImageJ Plugin.

### Statistical analyses

Statistical significance was performed using paired Student’s *t*-test. Differences were considered significant at *p* <0.05 and are indicated as follows: \**p* <0.05, \*\**p* <0.01 and \*\*\**p* <0.001.

## RESULTS

### Preparation of cells with labeled ORAI1

For the examination of the two-dimensional plasma membrane distribution of ORAI1 using LPEM, a HEK cell line was used pre-selected for a low background level of endogenous ORAI1. Details on this cell line, named CRI_1, are described elsewhere (9). For experiments with ORAI1 at rest, the cells were grown on microchips, as needed for LPEM (28), transfected with HA-tagged ORAI1, fixed, and labeled with QDs. Fig. 1a shows the side view of a scheme of QD655-labeled ORAI1 proteins, with all involved proteins and QDs drawn to scale. Note, that a QD may contain more conjugated streptavidin proteins than those shown in the scheme, but only one streptavidin per QD can bind to a biotinylated Fab. The HA-tag is located in the extracellular part of each of the six ORAI1 protein constituting the hexameric ORAI1 channel. In principle, each HA-tag can obtain a bound Anti-HA Fab. However, considering the size of the Fab and the narrow space between the 6 tags in the hexameric conformation, a 100% labeling efficiency is assumed to be impossible. Previously, we determined the labeling efficiency of these QDs for labeling HER2 proteins in whole cancer cells to amount to 27-83% depending on the exact conditions (27). A hexameric channel with two bound QDs is depicted in the scheme. The labels are placed in two possible spatial arrangements, the most upright position (shown with the two opaque labels), and the most laterally extended positions (shown with the transparent labels). Obviously, any position between these two extremes is also possible, whereby the distance between the center of a QD and the labeled ORAI1 protein is maximal 17 nm. In the STEM images, all labels are detected from a top view. The center position of a bound QD therefore indicates the presence of an underlying ORAI1 protein within a maximal radius of 17 nm, and a pair of QDs bound to the same ORAI1hexamer exhibits a maximal center-to-center distance of 35 nm.

Fig. 1b displays a typical light microscopy (LM) image from these cells in which several cells expressed ORAI1, which is visible from the GFP fluorescence co-expressed in a 1:1 molar ratio. The same cells displaying GFP fluorescence also show QD fluorescence (Fig. 1c), proving the specificity of the HA-ORAI1 tag. ORAI1 remained mostly homogenously distributed throughout the plasma membrane, which is expected for resting conditions (5).

To study the effect of SOCE-activation, the same cell line was used but here ORAI1-HA was co-expressed with STIM1 in a 1:3 ratio, and the cells were incubated for 15 min with 1 μM Tg at 37 °C for activation. The 1:3 ratio is known to be optimal for activation of the ORAI channels (29), and was achieved by transfected the expressing plasmids in this ratio (30). To examine the effects of a suboptimal STIM1:ORAI1 ratio, a third experimental group was also prepared in which ORAI1 and STIM1 were expressed in a 1:1 ratio, thus achieving a sub-maximal SOCE-activation after Tg incubation.

Control experiments were performed with another HEK cell line (CRI_STIM), lacking both endogenous STIM1 and STIM2 (16) to exclude that endogenous, pre-activated STIM1 or STIM2 had an influence on the distribution of ORAI1 (31) (see Fig. S2). A second series of control experiments were performed to examine possible influence of (residual) endogenous levels of ORAI1-3 proteins on the observed ORAI1 spatial distribution by expression in a triple ORAI1-3 knock-out cell line (CRI_2, see Fig. S3). Thirdly, to ensure that the observed effects after SOCE activation were not specific to or an artifact of Tg incubation, incubation with cyclopiazonic acid (CPA) was tested as well, as alternative SOCE activator (see Fig. S4).

### LPEM revealed ORAI1 clusters with supra-molecular dimensions

LPEM experiments of whole cells were carried out to map the positions of individual ORAI1 proteins for the CRI_1 cell line and QD655 labels. Using the QD fluorescence intensities of the LM images (Fig. 1c), cells were selected for analysis with STEM. In each experiment, cells from three groups were selected, exhibiting low-, medium, or high ORAI1 expression. Note that the average expression level differed between experiments, and, therefore, the relative expression levels in an experiment were used to define the three groups. STEM images revealing the positions of QD-labeled ORAI1 proteins were thus recorded from dozens of cells, yielding several hundred thousand of single ORAI1 positions (see Table 1). The selected cells covered the full range of ORAI1 expression levels found in our experiments. Fig. 2a depicts the typical ORAI1 distribution in a resting cell expressing ORAI1 at high levels, yielding a label density of 274/μm^2^, equivalent of the 90% percentile of the distribution of the label density in all recorded images. Shown here is the original STEM image with an overlay of automatically detected QD labels.

**TABLE 1.**
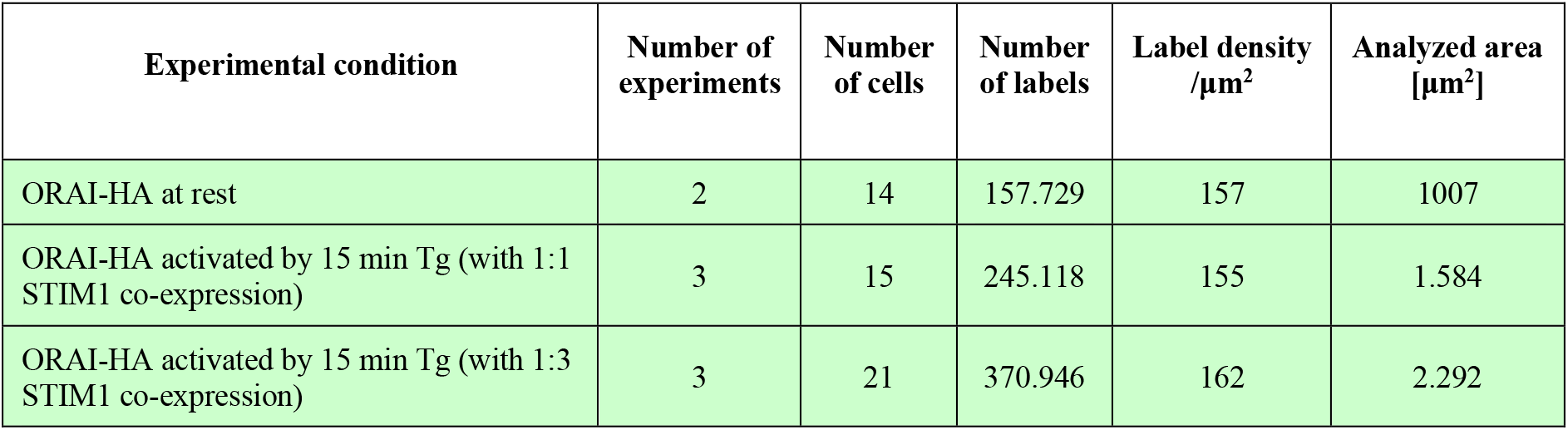
Data of ORAI1 labeling experiments including the experiment type. Conditions are at rest, at sub-maximal, and at maximal activation. Information provided is the number of experiments, the number of examined cells, the number of detected labels, the average label density, and the total analyzed membrane area.

**FIGURE 2.**
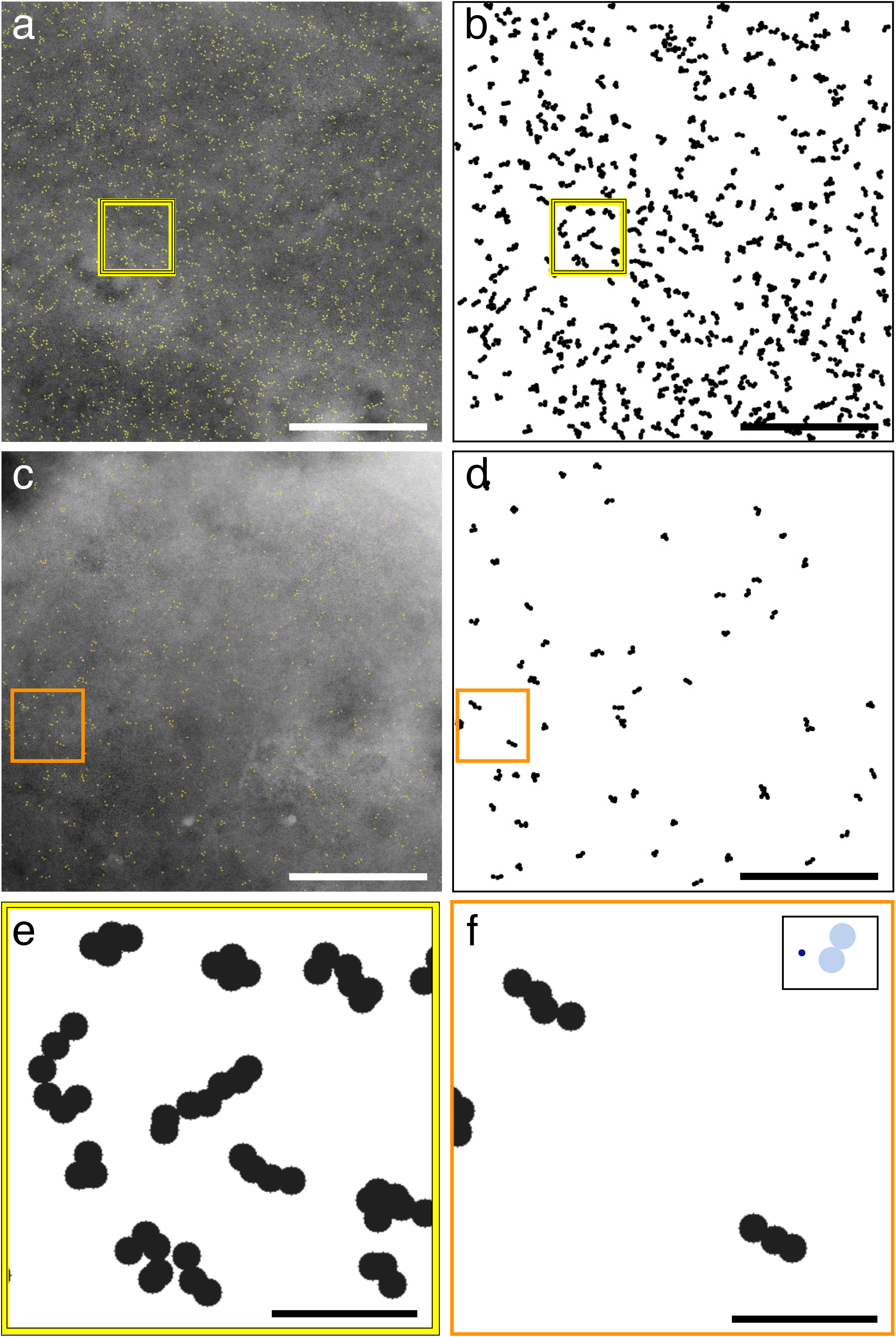
Scanning transmission electron microscopy (STEM) and cluster analysis of ORAI1 distributions in whole cells in liquid state. **(a)** Typical STEM image recorded from a resting cell with a relatively high expression of ORAI1 proteins. The electron-dense QD cores are outlined in yellow (n = 2.817 labels), revealing scattered, single labels as well as many ORAI1 clusters. **(b)** Result of the cluster analysis of the image shown in (a), excluding single labels and dimeric labels and showing only ORAI1 clusters consisting of 3 or more QDs. The labels are drawn as black circles (radius of 17 nm from the center position of each label), each indicating the possible spatial range of the underlying, QD-binding ORAI1 protein. The majority of the ORAI1 clusters display a linear or elongated shape, as can be seen in the examples in the boxed region, shown magnified in (e). **(c)** Exemplary image from a cell at rest with a low density in our study (n = 748 labels). **(d)** Cluster analysis showing all detected clusters of size ≥3 in (c). **(e)** and **(f**) show in more detail the elongated clusters in the boxed regions in (b), respect. (d). In the insert in **(f)** a dark blue circle depicts the dimension of a hexameric ORAI1 channel (7 nm extracellular diameter), two light blue circles of 35 nm diameter diameter) illustrate the space than can be maximally occupied by two labels bound to the same hexameric ORAI channel (see also Figure 1). The majority of the label clusters have major axis lengths exceeding the maximal distance of 70 nm, the limit for labels binding to the same ORAI1 hexamer, and thus indicate the presence of supramolecular assemblies of ORAI1 proteins. Scale bars in a-d = 1 μm, in e-f = 200 nm.

It appeared by eye that the overall ORAI1 distribution was homogenously distributed showing scattered labels throughout the whole imaged area. A more detailed view reveals that a large fraction of the labels showed distinct clustering in groups of less than ten labels. It has to be noted that even if all labeled target proteins would form oligomers, a fraction of mono-labeled oligomers will always appear, as well as a fraction of non-visible, non-labeled oligomers, because the labeling efficiency is usually below 100%. Generally, the share of under-labeled oligomers containing no or only one label depends on the number of oligomer-subunits and the labeling efficiency (32), which we assume to be ~30% for QD-labeled ORAI1, a value that was chosen based on results of our previous work including dimeric ORAI1 constructs (14). To enhance the visibility of ORAI distribution patterns, Fig. 2b shows the processed and cluster-analyzed image, where all QD-positions were marked by black circles of 35 nm diameter representing the area where the underlying, labeled ORAI1 protein resided, and only clusters containing more than 2 circles are shown. The processed image reveals that these clusters have a specific range of sizes and mostly elongated or linear shapes. Due to the high density of labels several clusters appear so close that they touch each other. This close proximity hampered the measurement of their spatial dimensions and shapes. In low ORAI1 expressing cells, the lower density of expressed ORAI1 was expected to lead to more accurate determination of these clusters, due to less frequent touching of cluster boundaries. Therefore, a data set from low expressing cells, was analyzed separately. An example of such an image, with a label density of 74/μm^2^, equivalent of the 18% percentile of the label density is shown in Fig. 2c. The corresponding processed and cluster-analyzed version is shown in Fig. 2d. From visual inspection of the image it appears that clusters >2 mostly exhibit an elongated shape. To compare high-expressing cells with low expressing cells, an example of each group is displayed magnified in Fig. 2e and f, respectively. Due to less coalescing clusters in low expressing cells they can be better discerned and more accurately measured; two examples can be discerned in Fig. 2f.

To compare the label and cluster dimensions with the dimensions of the underlying, invisible, ORAI1 protein hexamers, the insert in Fig. 2f shows a small circle of 7 nm diameter, representing the extracellular part of a hexameric ORAI1 channel, and two circles of 35 nm diameter indicating the maximal distance that can be occupied by two labels bound with the most extended configuration to the same ORAI1 channel (see Fig. S1). One can see that most clusters exceed this dimension, and can thus not have originated from single ORAI1 channels but must have resulted from underlying clustered channels. Because the membrane proteins had been immediately fixed prior labeling, it can be reasonably assumed that the clusters larger than 70 nm did not originate from labeled ORAI1 channels that disassembled after labeling, and then diffused in the plasma membrane leading to a larger distance. Moreover, a number of clusters contained more than 6 labels, and many of these clusters formed chains up to 120 nm length.

To characterize the dimensions of these chain-like protein arrangements, they were examined for their maximum dimension, by measuring the length of their major axis, and for their aspect ratio. These measurements were automatically performed on the processed images showing each label as a 35 nm circle. Touching and coalescing groups of labels were classified as clusters, whereby only clusters containing >2 labels were analyzed.

An important question is if the observed elongated clusters were the result of random orientation or reflected an underlying organization of ORAI1 hexamers. To answer this question, images were analyzed of simulated random distributions of labeled hexameric ORAI1 channels. The simulated images were created from sets of individual STEM images recorded from low expressing cells. Label distributions were simulated by individually matching the label density of each image, assuming randomly distributed hexameric ORAI1 channels and a labeling efficiency of 30%. When cluster >2 from QD655 data of low expressing CRI_1 cells at rest were compared with their matching simulation, they were 19% longer and had 24% larger AR than those from the simulations (see Fig. S6) (both parameters: *p* <0.001). A similar comparison of cluster from data from high expressing cells with those found in matching simulations confirmed the differences found in low expressing cells. Here the values from the STEM images found for the long dimension, and for the AR were 13%, respect. 15% higher than those from the simulations (see Fig. S6) (for both parameters: *p* <0.001). The relatively lower difference between data and simulations in the high expression data, compared to the low expression data, is caused by a partially biased cluster detection through a rise in the fraction of clusters touching each other. A more detailed comparison between the results from low and high expressing cells at rest is provided in the following section (see below).

To examine a possible influence of STIM and ORAI3 proteins on the ORAI1 cluster formation similar control experiments were performed in cells lacking endogenous STIM (CRI_STIM cell line) (Fig. S2) and in triple ORAI1-3 knock-out cells (CRI_2 cell line, Fig. S3). Cluster analysis from both cell lines revealed similarly elongated cluster >2 and a similar fraction of ¾ exceeding the limit of 70 nm, thus matching the findings found in the cells of the mainly used CRI_1 cell line. We can thus exclude that the formation of these supramolecular ORAI1 complexes depends on STIM or ORAI3 proteins.

To examine an effect of the applied QD label on the results from the cluster analysis, similar experiments and analysis of the recorded STEM images were performed using the smaller QD565 as ORAI1 labels (see Fig. S6). Comparison of cluster results from STEM data sets of QD565 labeled ORAI1 in resting cells, with a low expression level, with their matching simulations, were performed with the main cell line CRI_1 as well as with the CRI_2 cell line. When compared to each other, the two cell lines showed no difference (*p* = 0.85 for the length, respect. *p* = 0.14 for AR). In contrast, comparing the cluster from QD565 STEM images with their matching simulation, showed that the cluster in the images were by 21% (CRI_1), respect. 18% (CRI_2) longer, and had a 26% higher AR (for both cell lines) than cluster in the matching simulations (all *p*<0.0001). These relative differences between cluster >2 in STEM images and simulations match the 19%, respect. 24% for the relative difference in dimension, respect. AR, found with the QD655 labeling (see section above). It can thus be concluded that the supra-molecularly sized ORAI1 clusters were neither randomly appearing structures, nor influenced by the type of QD label, or the cell line used, but must have originated from a biological interaction leading to the spatial organization of ORAI1 hexamers in the plasma membrane.

### SOCE activation partially relocates ORAI1 into distinct accumulation areas

After maximal SOCE-activation, typical bright ORAI1 punctae were seen in the LM images (Fig. 3a). LPEM revealed that ORAI1 accumulated in oval areas, presumably ER plasma membrane contact zones. These regions of condensed ORAI1 accumulations had diameters of 1.3 ± 0.5 μm for the major axis, 0.7 ± 0.2 μm for the short axis, an aspect ratio of 2.0 ± 0.6, and an area size of 0.9 ± 0.5 μm^2^ as directly measured from STEM images (see Fig. S7), matching the size of punctae reported earlier (2). Similar ORAI1 punctae were also found after SOCE-activation with CPA, although with on average smaller spatial dimensions and aspect ratios (see Fig. S4). Fig. 3b displays the distribution of all clusters >2 found in a typical STEM image from a maximally SOCE-activated cell. ORAI1 distributions in the punctae were crowded (shown magnified in Fig. 3c), but it is still possible to regionally recognize elongated clusters, suggesting that the dense ORAI1 accumulation areas included chain-like supra-molecular ORAI1 clusters. Outside the punctae regions, the spatial distribution of ORAI1 clusters >2 (shown magnified in Fig. 3d) resembled those found in the resting cells.

**FIGURE 3.**
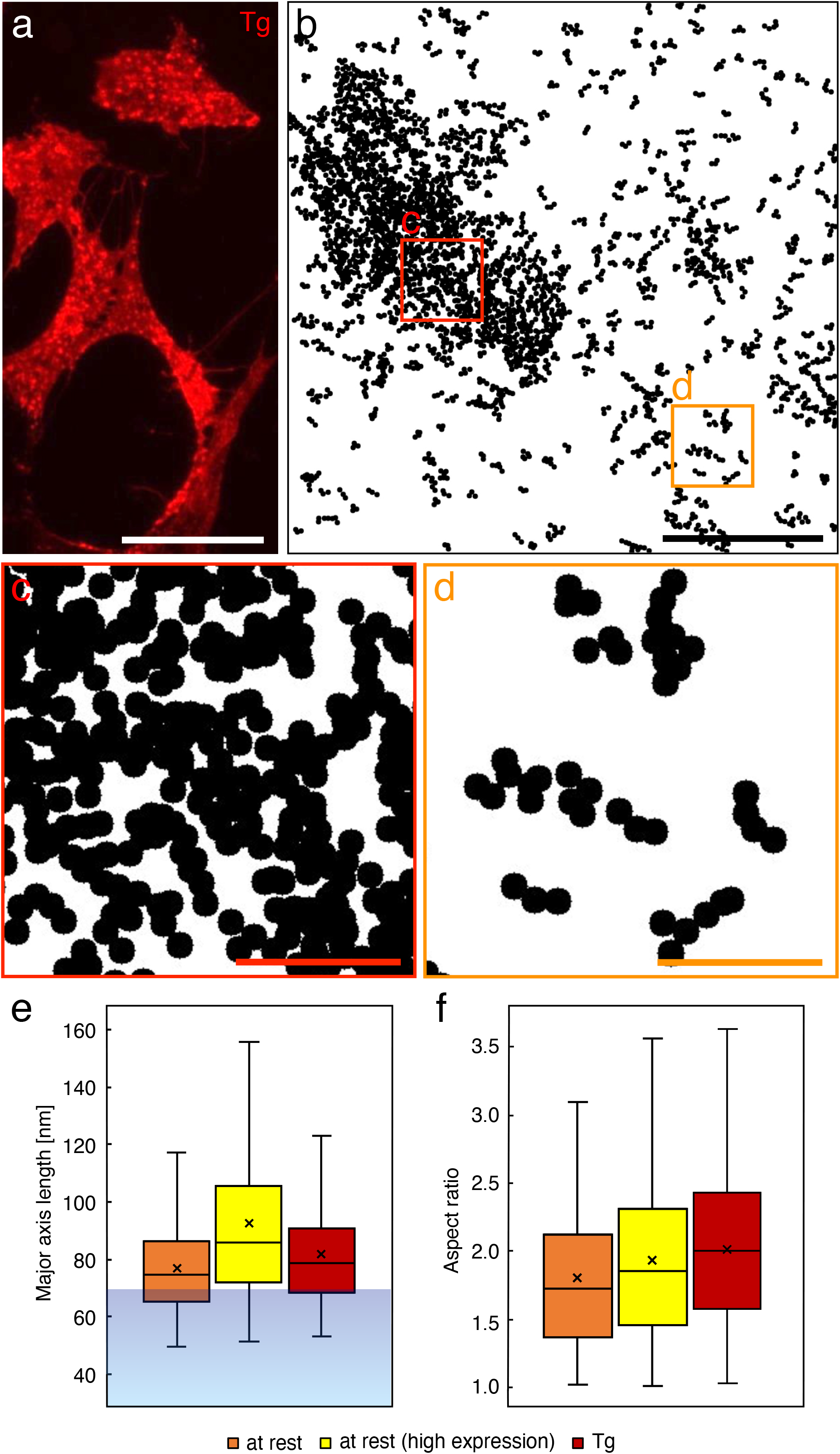
ORAI1 distributions 15 min after Thapsigargin (Tg) in maximally SOCE-activated cells (expressing ORAI1 and STIM1 in a 1:3 ratio). **(a)** Fluorescence microscopy image showing the QD signal. The in-homogenous distribution of QD fluorescence into bright circles indicates the assembly of ORAI1 in punctae. **(b)** Cluster analysis of a STEM image recorded after maximal SOCE-activation, showing all clusters of size >3. A densely ORAI1 filled, ellipsoid area is visible on the left side, surrounded by more loosely scattered, elongated clusters. **(c)** and **(d**) show the boxed regions in (b) in more detail. The clusters found within the dense region (c) are so condensed that their elongated shape can only partially be recognized (mainly in the lower left area). (d) show the cluster distribution outside of the dense region where the high aspect ratios of the clusters emerge more clearly. **(e)** Quantitative results showing box plots for the measured length of the major axis length of all entirely displayed clusters of size >2, detected in STEM images from cells at rest showing low expression (orange) (n = 436 clusters, average density = 62 labels/μm^2^), high expression (yellow) (n = 1.548 clusters, average density = 242 labels/μm^2^), and from regions outside the punctae in maximally Tg activated cells with low expression (n = 277 clusters, average density = 53 labels/μm^2^). The corresponding percentile values of the distribution of the label density, were 13%, respect. 84%, and 8%. In all three experimental groups, the vast majority of the clusters are longer than 70 nm, which is the max. range for QDs attached to the same ORAI1 hexamer (represented by the transparent blue region <70 nm in the plot). **(f)** Determination of the aspect ratio (AR) for all clusters of size >2, reveals similar values in resting low and high expressing cells and in max. Tg activated cells outside the punctae. Scale bar in a = 50 μm, in b = 1 μm, and in c and d = 200 nm.

Fig. 3e and f show the results of quantitative analysis of clusters >2 data sets for their maximal dimension, and their aspect ratio, derived from cells at rest, of low- and high expressing cells-, and from low expressing cells after max. SOCE-activation, measured outside of the punctae regions. For this latter group the STEM images were chosen such, that they contained no or just a few small punctae. The results in Fig. 3e show that, independent of the activation status or the expression level, about ¾ of all detected cluster >2 exceed 70 nm, reaching up to 120 nm of length, and thus indicate underlying supramolecular arrangements of ORAI1. Note that at rest, high ORAI1 expressing cells seem to have a 14% larger cluster dimensions than low expressing cells, whereas in max. activated cells with low ORAI1 expression, cluster outside the punctae are only 4% larger (*p* <0.001 between all groups). However, larger cluster dimensions in the high expressing cells are simply due to more touching and coalescing clusters in high expressing cell, biasing the automated measurement, which considers several touching cluster as a single cluster unit. The results of cluster AR measurement, shown in Fig. 3f, reveal a marginal increase of 4% in high expressing versus low expressing cells at rest, but a 14% higher AR after activation, possibly reflecting a stretching effect, for instance when clusters are drawn towards the punctae. For all comparisons in the quantitative cluster measurements of Fig. 3e and f *p*<0.001.

### ORAI1 patterns under limiting STIM conditions

To better examine the ORAI1 distribution in the punctae, we examined cells after sub-maximal SOCE-activation, caused by insufficient abundance of STIM1 proteins, after a 1:1 co-expression ratio with ORAI1. In these cells, a somewhat different clustering pattern of ORAI1 emerged compared to the maximal SOCE activation. Accumulated ORAI1 appeared in larger, irregularly shaped, and more diffuse patches (Fig. 4a). In the STEM images, the borders of ORAI1 accumulation areas emerged less defined than in maximally activated cells, this effect is visible in Fig. 4b, showing the processed and cluster-analyzed version of a typical STEM image from cells with a lowered STIM:ORAI ratio after 15 min of Tg incubation. The right half of the image shows accumulating ORAI1 clusters >2, however, the degree of label crowding was less dense, compared to punctae regions in maximally activated cells (Fig. 3b). Magnifications of the marked regions in the punctae region in Fig. 4b are shown in Fig. 4c, and a region outside is shown in Fig. 4d. As for the resting cells and the maximally activated cells, also the sub-maximally activated cells showed elongated ORAI1 clusters outside and clearly also inside the accumulation areas. After submaximal activation the free space between the cluster was larger than after maximal activation, better disclosing that small groups of two or three chain-like ORAI1 clusters had aligned in a parallel manner (Fig. 4c, and Fig. S8). These results support the concept that upon activation, dimers of activated STIM1 proteins crosslink neighboring ORAI channels (16), which in this case appear to be pre-arranged as linear complexes. Fig. 4e shows that the clusters >2 found outside from punctae in sub-maximally activated cells (measured from high expressing cells) were the same as those found in resting cells.

**FIGURE 4.**
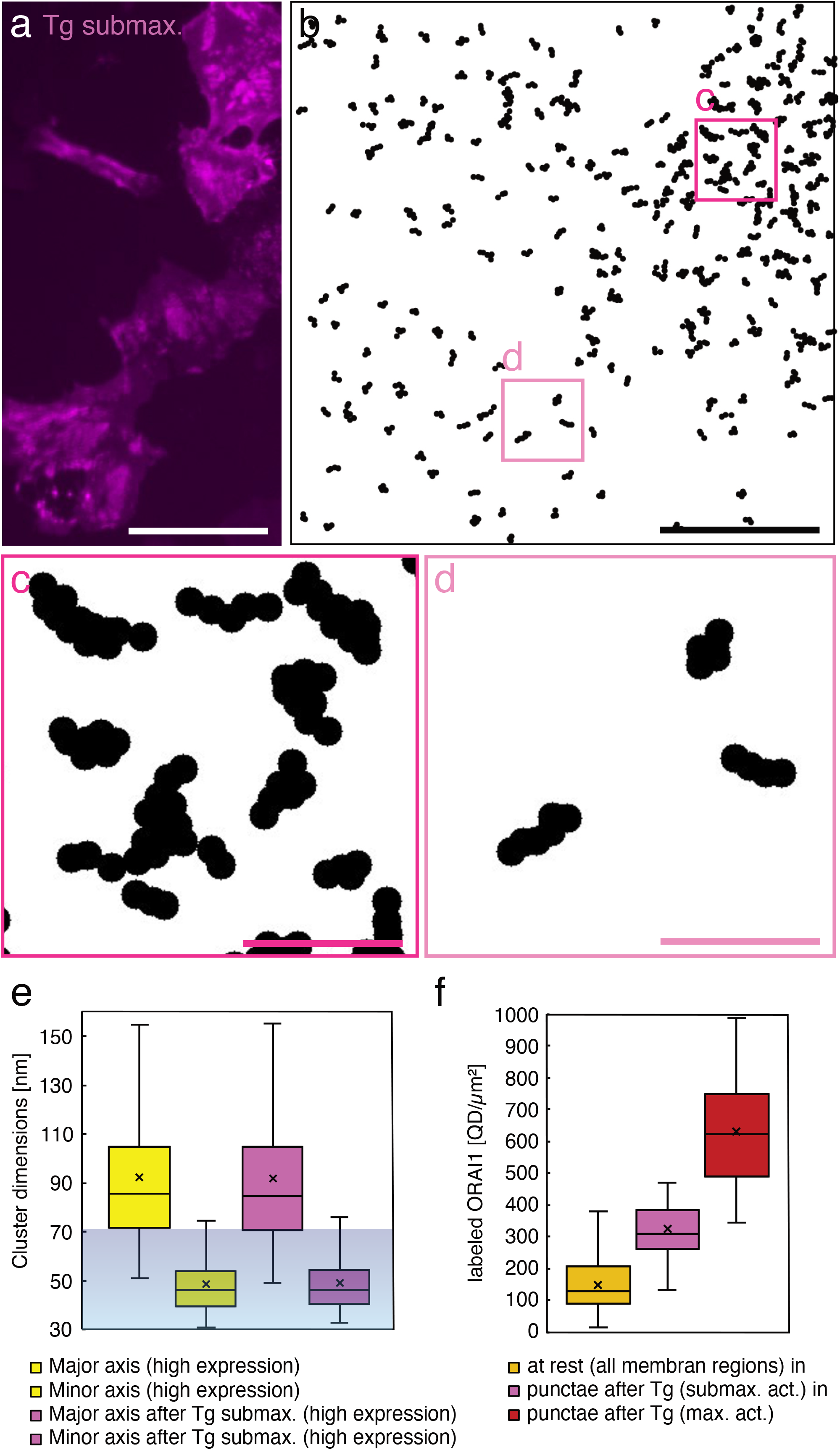
ORAI1 distributions 15 min after Thapsigargin (Tg) in submaximally SOCE-activated cells (expressing ORAI1 and STIM1 in a 1:1 ratio). **(a)** Fluorescence microscopy image of the QD signal showing accumulations of ORAI1 in bright patches. Most of these regions with denser ORAI1 distributions are larger and blurrier, than the punctae found after maximal SOCE-activation (compare to Figure 3a). **(b)** Cluster analysis of an exemplary STEM image recorded after submaximal SOCE-activation, showing all clusters of size >2. The clusters are scattered throughout the image area, except for the area in the right upper part of the image, were clusters have accumulated. Most clusters have similar elongated shapes those found in resting and in maximally SOCE-activated cells (compare to Figure 2e and f, and Figure 3d). **(c)** and **(d)** show magnified details of the boxed regions in (b), revealing that several clusters in the condensed puncta area have aligned during the redistribution process elicited by the SOCE-activation. Clusters found outside the condensed area (d) remained mostly separated. **(e)** Spatial dimensions of label clusters >2 detected in STEM images from cells at rest (with high ORAI1 expression) (yellow, n=1548), and from cells after submaximal SOCE-activation (pink, n = 1344). 75% of the clusters in both groups have major axis lengths exceeding the spatial limit for labels bound to the same ORAI1 hexamer, whereas the length of the minor axis remains almost always below this limit. The dimensions of clusters from cells at rest and after activation are not significantly different. **(f)** Comparison of QD-labeled ORAI1 densities in STEM images recorded from all plasma membrane regions of resting cells (dark yellow, n = 113 STEM images), with densities in punctae from sub-maximally SOCE-activated cells (pink, n = 36 punctae), and punctae from maximally SOCE-activated cells (red, n = 10 punctae). In cells with similar average label densities, submaximal SOCE-activation had doubled the regional density of labeled ORAI1 in punctae, whereas maximal activation resulted in a four-fold increase in density. Scale bars in a and b = 50 μm, in c and d = 200 nm.

Fig. 4f presents the results of a quantitative analysis of the densities of labeled ORAI1 in resting cells and after activation in punctae. Analyzed were images from selected cells displaying similar average densities of 166±80/μm^2^, for sub-maximally activated cells, respect. 172±55/μm^2^, for maximally activated cells. Submaximal activation approximately doubled the average label density from 155 ±87/μm^2^ (yellow), as measured on membrane areas from all resting cells, to 334 ±108/μm^2^ inside regions of punctae, and maximal activation had increased the density inside punctae to 640 ±193/μm^2^.

To estimate the percentage of ORAI1 proteins relocating into punctae areas after max. SOCE-activation, 44 STEM images (from 11 cells) with fully depicted punctae areas were analyzed (the remaining 125 STEM images contained only partially depicted or no punctae). In the selected images the average density of labeled ORAI1 per cell matched the density found in all images from this experimental group of (see Table 1 and 2). The label densities inside and outside the punctae areas, and the size of the punctae area were determined, first from every image and then per cell, for the ratio of label density inside/outside punctae. This ratio was found to be 3.7 ±1, the fraction of plasma membrane covered by punctae amounted to 14%. It should be noted however, that the excluded STEM images contained also images without punctae, therefore, a value 14% is slightly overestimated. The true value should lie between these 14% and a minimum of 5%, which would apply if only the fully depicted punctae were present. This proposed range of 5 – 14% fits well the earlier reported share of 2 – 13% plasma membrane regions with underlying cortical ER (34). For calculating the percentage of relocated ORAI1 into punctae the value of 3.7, representing the average ratio for label density outside/inside punctae, was applied to the 5 - 14% share, resulting in 14 - 24% of all ORAI1 proteins redistributed into punctae after maximal activation. Insertion of additional ORAI1 proteins from intracellular stores into the regions of punctae (12) does not seem likely since the average surface density of ORAI1 was similar between resting- and activated cells (see Table 1), and the density of ORAI1 dropped by ~10% in the areas outside of the punctae compared to the overall density in cells at rest. Therefore, it seems rather likely that ORAI1 predominantly relocates from the surrounding plasma membrane into punctae.

**TABLE 2.** Measurements of ORAI1 distributions inside and outside punctae after maximally store-operated calcium entry (SOCE) activated cells.

| Experimental condition | Number of experiments | Number of cells | Number of labels | Analyzed area [$\mu\text{m}^2$] | % area occupied by punctae | label density inside punctae /$\mu\text{m}^2$ | label density outside punctae /$\mu\text{m}^2$ |
| --- | --- | --- | --- | --- | --- | --- | --- |
| ORAI-HA activated by 15 min Tg (with 1:3 STIM1 co-expression) | 2 | 11 | 143.806 | 510 | 14 | 694 | 139 |

**TABLE 3.** List of control experiments performed with other cell lines, other labels and other methods of SOCE-activation.

| Experimental condition | Number of experiments | Number of cells | Number of labels | Label density /$\mu\text{m}^2$ | Analyzed area [$\mu\text{m}^2$] |
| --- | --- | --- | --- | --- | --- |
| ORAI-HA in CRI_STIM cells, activated by 15 min Tg | 2 | 8 | 80.887 | 120 | 672 |
| ORAI-HA activated by 15 min CPA (with 1:3 STIM1 co-expression) | 1 | 8 | 153.102 | 203 | 753 |
| ORAI-HA at rest, labeled with QD565 | 1 | 11 | 76.126 | 304 | 251 |
| ORAI-HA in CRI_2 cells at rest | 2 | 11 | 91.548 | 96 | 953 |
| ORAI-HA in CRI_2 cells at rest, labeled with QD565 | 4 | 14 | 81.303 | 277 | 294 |

### Punctae are not the only type of structures in activated cells

Besides the typical ellipsoid ORAI1 punctae found after SOCE-activation several other twodimensional (2D) structures were found as well in the plasma membrane. Fig. 5a shows examples of micron-long, parallel running ORAI1 formations (indicated with transparent red lines). These ORAI1 strand-like patterns meandered in regions corresponding to areas with a brighter background than the surrounding. Since the contrast in STEM increases with mass, the brighter shapes represent thickened cellular regions. In addition, four oval ORAI1 punctae were present. Fig. 5b is the label-to-circle processed image recorded from a cell with a low ORAI1 expression, also in this case several micron-long strands can be discerned that run in parallel. Thus, rules out that such 2D-structures were an artifact of highly overexpressed ORAI1. They were found in 27% of the recorded STEM images from maximally SOCE-activated cells. Fig. 5c, shows examples of ORAI1 punctae differing from the typical ones by their ring-like structures; the largest ring-structure had densely accumulated strands of ORAI1 in its center. These peculiar 2D-structures must have been created by an underlying mechanism elicited by SOCE-activation, and were also found in experiments using CPA induction of SOCE-activation (see Fig. S4).

**FIGURE 5.**
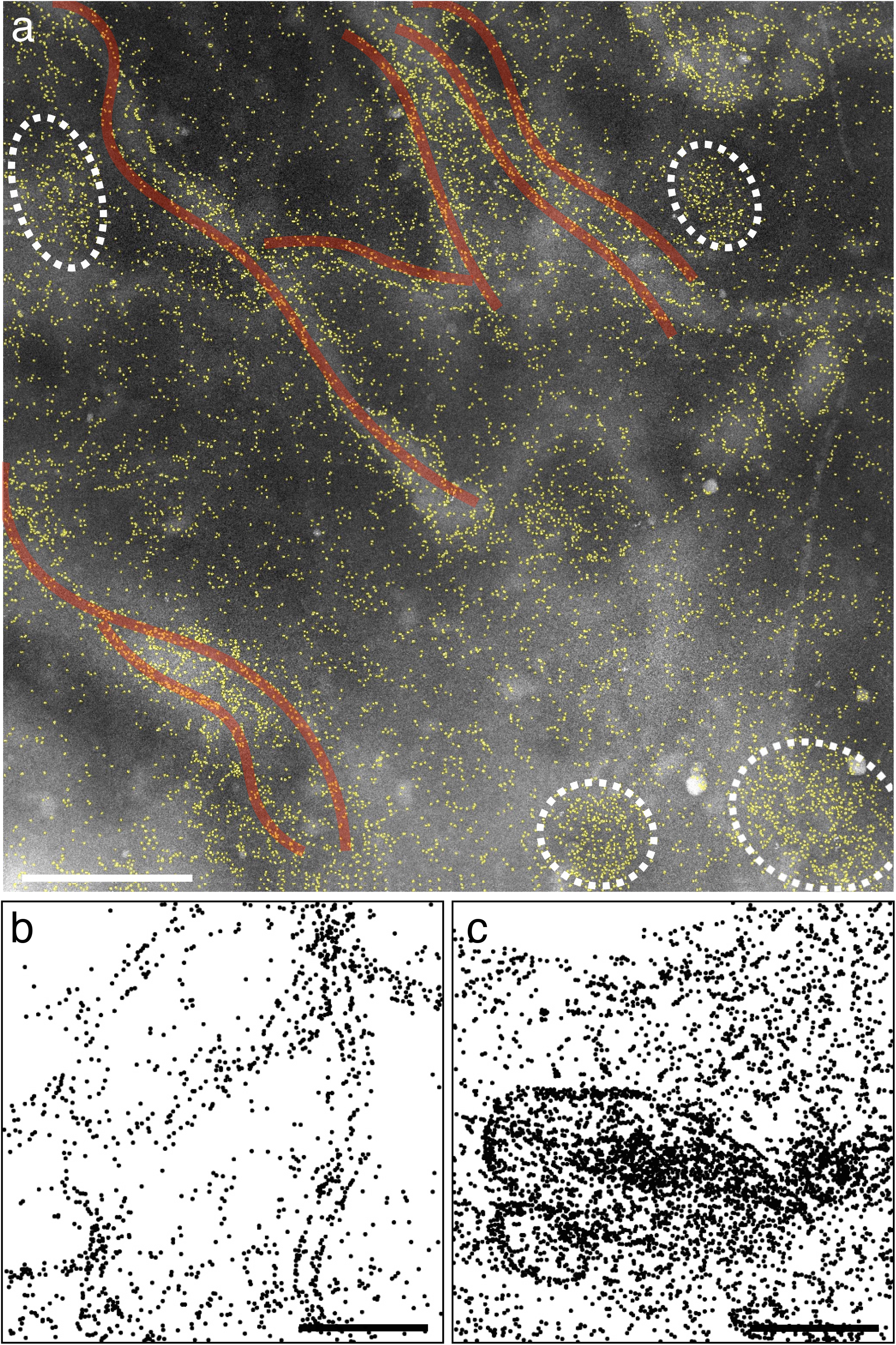
ORAI1 distributions in activated cells display a variety of 2D structures. **(a)** Original STEM image (with QD labels outlined in yellow) gives an impression of the regional cellular morphology, with thicker, cellular areas appearing brighter than thinner ones. A bulging membrane topography often matches with dense accumulations of ORAI1. Typical, ellipsoid punctae (outlined by dashed lines) and nearby running, several micrometer-long strands of aligned ORAI1 (transparent red traces) can be seen. **(b)** and **(c)** show other processed STEM images from cells with a relatively lower (b), respect. higher (c) ORAI1 expression level, displaying all detected, labeled ORAI1, including monodisperse and dimeric label conformations, as 35 nm diameter circles. In (b), micrometer-long chains of ORAI1 formations emerge, several of them running in parallel. In (c), ellipsoid, ring-like structures of ORAI1distributions of different dimensions are visible, the larger of these structures encloses a central region of condensed ORAI1. Scale bars = 1 μm.

## DISCUSSION

Our study aimed to elucidate how the spatial organization of ORAI1 proteins in the plasma membrane changes upon Ca^2+^ channel activation. We have used an HA-tagged ORAI1 protein in combination with a two-step QD labeling approach, resulting in a label:target ratio of <1, thereby excluding any clustering of more than one QD at a single ORAI1 protein (9). It can also be excluded that the applied QD-labeling protocol induces artificial clustering of membrane proteins. This proof was published for a similar QD-labeling approach of another membrane protein, namely HER2, where randomly distributed QD label positions occurred in a phenotypic distinct subpopulation of HER2 overexpressing cancer cells (26, 35), and that finding was later confirmed in a fraction of human cancer cells from biopsies (36).The proteins were detected in the intact and hydrated cell, using correlative light microscopy and LPEM, yielding the position information of several hundred thousand of single-labeled ORAI1 proteins.

In resting cells, we discovered that ORAI1 often assembled in chain-like structures extending up to 120 – 130 nm. ORAI1 proteins thus generally tend to form supra-molecular arrangements. Considering a predominant distance of ~15 nm between adjacent ORAI1 channels in punctae, as measured by electron microscopy (37), the underlying chain-like conformation could contain between 2 and 8 ORAI1 channels. The existence of such multichannel ORAI1 conformations contradicts the assumption that ORAI1 channels predominantly exist as independent entities, randomly distributed in the plasma membrane (30, 37), the authors further suggested that single channels were freely diffusing within these patches. Although they also used EM to examine overexpressed ORAI1, the proteins were imaged without labels, and in freeze-fractured cells (37), posing challenges for the discrimination of ORAI against all other, neighboring membrane proteins. Nevertheless, the authors stated the occasional detection of ORAI1 channels in specific arrangements of pairs and short chains.

In the STEM images recorded after SOCE-activationORAI1 clusters had been trapped into the regions of punctae where, depending on the availability of STIM proteins, they often by aligned side-by-side, as found after submaximal activation, or condensed further until reaching a rather confluent pattern, after max. activation. Measurements of ORAI1 densities in maximally activated cells, inside and outside punctae, revealed on average a 4-fold increase within the regions of punctae, achieved by the redistribution of a minor ORAI1 protein fraction, ranging between 14 and 24% of all plasma membrane-bound ORAI1. Our results do not support earlier findings reporting the insertion of ORAI1 from intracellular pools into punctae after SOCE-activation (12). Such a replenishment of plasma membrane localized ORAI1 would lead to an increase in the average ORAI1 surface density in SOCE-activated cells versus cells at rest, however the average ORAI1 densities in our experiments remained similar. A possible reason for this discrepancy might be the exclusive use of formaldehyde as fixation agent in this earlier study, which is known to insufficiently fix membrane proteins (38, 39), and to induce artificial redistributions of ORAI1 proteins (40).

In resting cells, Perni et al reported micron-sized patches with ragged edges, assuming these areas contained dense accumulations of ORAI1 channels (37). We could not confirm this finding, instead, labeled ORAI1 distributed rather evenly throughout the plasma membrane of resting cells, with mono-dispersed and clustered label formations. This discrepancy might lie in differences between the used preparation and imaging techniques. Our comparison between the cluster analysis of STEM image data with their density-matching simulations, led us to propose that a significant fraction of the ORAI channels formed linear arrangements. For all examined experimental groups, the dimensions and aspect ratios of labeled-ORAI1 clusters >2 in the STEM images were significantly larger (p < 0.001) than those of clusters >2 found in their matching simulations (see Fig. S5 and Fig. S6). High-order aggregates of non-solubilized ORAI proteins were found earlier with the biochemical technique of Native Gel Electrophoresis using perfluorooctanoic acid (PFO), but these results were interpreted as technical artifacts (5). Also our recently published results from biochemical studies with Blue Native Polyacrylamid Gel Electrophoresis revealed that a significant fraction of ORAI1 assembled in larger order oligomers than dimers, mainly appearing as a complex with an apparent molecular mass in the range of the calculated mass of two hexamers (9). Furthermore, ORAI1 diffusion coefficients were reported to exhibit a large range of different values, whereby most ORAI1 proteins had sub-diffusive velocities (2), possibly caused by differently sized molecular ORAI1 assemblies (41), a finding which fits to the idea of supra-molecular ORAI1 channel arrangements.

Concerning the larger, peculiar 2-D assembly patterns detected in SOCE-activated cells, such as micrometer-long strands and ring-like structures, we were unable to find matching reports in the literature. Perni et al did confirm the presence of dense ORAI accumulations in puncta after SOCE-activation, but their observations did not include strand- and ring-like 2D structures. This discrepancy is likely due to the small width of these strands, often of a single labeled ORAI1, and the associated difficulty of discerning such small features amongst the other neighboring proteins in the crowded environment of the plasma membrane without using labels.

A possible function of clustering of ORAI1 in supramolecular arrangements could be a more efficient concentration process of ORAI1 Ca^2+^ channels in punctae upon SOCE activation compared to a mechanism via the collection of single channels. The capturing and dragging of only one ORAI1 protein belonging to a supramolecular cluster will drag all the others with it, until the whole cluster reaches the ER plasma membrane contact zone. A confluent filling of the plasma membrane areas, located above accumulated STIM in the ER plasma membrane contact zones, by supra-molecular ORAI1 clusters, instead of single ORAI1 channels, would not only be faster, but would also help in achieving a more homogenous filling, avoiding a jamming of ORAI1 channels at the periphery of the punctae, which would hinder and delay the filling of the more central areas. An amplifying effect, due to the import of supra-molecular ORAI clusters instead of single ORAI1 channels, would lower the required amount of effectively trapped and transported ORAI1 far below the 14 to 24% of ORAI1 proteins found to be redistributed after maximal SOCE-activation. Such an amplifying effect would, for instance, play an important role in the immune system, by accelerating the formation of immune synapses between T cells and antigen-presenting cells (42).

The comparison of ORAI1 distributions in accumulation zones in sub-maximally activated cells revealed less dense accumulations of ORAI1 clusters than in maximally activated cells. Yet, often the accumulation areas showed aligned supra-molecular ORAI1 clusters, further supporting the concept that the import of preformed, supra-molecular ORAI1 clusters contributed to the accumulation of ORAI1 in punctae.

The existence of supra-molecular protein clusters in the plasma membrane, with similar dimensions and numbers of involved proteins as detected in this study for ORAI1 proteins possibly points to a general organization principle in the plasma membrane, as such protein arrangements were also found for other membrane proteins, mainly by using superresolution fluorescence microscopy. After over-expression in HEK cells, for instance, about half of the NMDA receptors were found in clusters containing up to 12 receptors (43). Also, for the family of G-protein coupled receptors, about half of the μ-opioid receptors, and 85% of κ-opioid receptors were reported to reside in clusters with dimensions of 80 – 100 nm, comprising on average 8 and 9 receptors, respectively (44). Assemblies of 10 or more receptors were also reported for the epidermal growth factor receptor (EGFR) (45), and chains containing six or more receptors for the closely related human epidermal growth factor receptor 2 (HER2) (46).

It is not known if a linking protein, responsible for the organization of supra-molecular ORAI1 clusters, exists. STIM proteins fulfill such an ORAI1 binding function and concomitant to binding, activate ORAI1 channels in the punctae. But, outside these regions and at rest, clustering of ORAI1 due to STIM can be excluded, as shown by control experiments (see Fig. S2). The possible mechanism behind the clustering of ORAI1 could involve homo-oligomerization (47, 48), the involvement of cytoskeletal proteins (49, 50), or septins, recently shown to be involved in cellular calcium signaling through ORAI1 (41), as well as a range of other known ORAI1-binding proteins (51, 52).

In conclusion, we found that most ORAI1 channels were organized in small, elongated clusters of supra-molecular size ranging from 50 – 130 nm. After SOCE-activation, 14 to 24% of plasma membrane-bound ORAI1 reorganized into regions (punctae) of high surface density of oval shape. In addition, a variety of other 2D structures appeared, most of them micron-long strands, and also ring-like patterns. We propose that the supra-molecular ORAI clusters serve an amplifying function for SOCE-activation, by reducing the required number of directly trapped ORAI1 proteins for subsequent accumulation in punctae, as needed for fast and efficient activation by STIM1 proteins in order to induce Ca^2+^ influx.

## SUPPORTING MATERIAL

Supporting Material can be found online at

## AUTHOR CONTRIBUTIONS

D.B.P, B.A.N, and N.J designed research; D.A. provided ORAI constructs and cells; D.B.P. performed experiments, and analyzed data; D.G. created ImageJ Plugins for image processing and data analysis, as well as the simulation algorithm, D.B.P., B.A.N. and N.J. wrote the paper.

## ACKNOWLEDGEMENTS

We thank Sercan Keskin, and Peter Kunnas for help with the STEM, Tabea Trampert for editing the figures, Don Gill for the kind gift of CRI_STIM cells, and E. Arzt for his support through INM. The research was funded by the DFG SFB1027 (project C7).

## REFERENCES

1. Dynes, J.L., A. Amcheslavsky, and M. D. Cahalan. 2016. Genetically targeted single-channel optical recording reveals multiple Orai1 gating states and oscillations in calcium influx. Proc. Natl. Acad. Sci. 113:440–445.

2. Wu, M. M., E. D. Covington, and R. S. Lewis. 2014. Single-molecule analysis of diffusion and trapping of STIM1 and Orai1 at endoplasmic reticulum–plasma membrane junctions. Mol. Biol. Cell 25:3672–3685.

3. Madl, J., J. Weghuber, R. Fritsch, I. Derler, M. Fahrner, I. Frischauf, B. Lackner, C. Romanin, and G. J. Schutz. 2010. Resting State Orai1 Diffuses as Homotetramer in the Plasma Membrane of Live Mammalian Cells. J. Biol. Chem. 285:41135–41142.

4. Maruyama, Y., T. Ogura, K. Mio, K. Kato, T. Kaneko, S. Kiyonaka, Y. Mori, and C. Sato. 2009. Tetrameric Orai1 Is a Teardrop-shaped Molecule with a Long, Tapered Cytoplasmic Domain. J. Biol. Chem. 284:13676–13685.

5. Penna, A., A. Demuro, A. V. Yeromin, S. Y. L. Zhang, O. Safrina, I. Parker, and M.D. Cahalan. 2008. The CRAC channel consists of a tetramer formed by Stim-induced dimerization of Orai dimers. Nature 456:116–U112.

6. Mignen, O., J. L. Thompson, and T. J. Shuttleworth. 2008. Orai1 subunit stoichiometry of the mammalian CRAC channel pore. J. Physiol. 586:419–425.

7. Hou, X., L. Pedi, M. M. Diver, and S. B. Long. 2012. Crystal Structure of the Calcium Release–Activated Calcium Channel Orai. Science 338:1308.

8. Cai, X., Y. Zhou, R. M. Nwokonko, N. A. Loktionova, X. Wang, P. Xin, M. Trebak, Y. Wang, and D. L. Gill. 2016. The Orai1 Store-operated Calcium Channel Functions as a Hexamer. J. Biol. Chem. 291:25764–25775.

9. Alansary, D., D. B. Peckys, B. A. Niemeyer, and N. de Jonge. 2020. Detecting single ORAI1 proteins within the plasma membrane reveals higher order channel complexes. J. Cell Sci. 133.

10. Liou, J., M. L. Kim, W. D. Heo, J. T. Jones, J. W. Myers, J. E. Ferrell, and T. Meyer. 2005. STIM is a Ca2+ sensor essential for Ca2+-store-depletion-triggered Ca2+ influx. Curr. Biol. 15:1235–1241.

11. Mercer, J. C., W. I. DeHaven, J. T. Smyth, B. Wedel, R. R. Boyles, G. S. Bird, and J. W. Putney. 2006. Large store-operated calcium selective currents due to co-expression of Orai1 or Orai2 with the intracellular calcium sensor, Stim1. J. Biol. Chem. 281:24979–24990.

12. Hodeify, R., S. Selvaraj, J. Wen, A. Arredouani, S. Hubrack, M. Dib, S. N. Al-Thani, T. McGraw, and K. Machaca. 2015. A STIM1-dependent ‘trafficking trap’mechanism regulates Orai1 plasma membrane residence and Ca2+ influx levels. J. Cell Sci. 128:3143–3154.

13. Peckys, D. B., D. Alansary, B. A. Niemeyer, and N. de Jonge. 2016. Visualizing Quantum Dot Labeled ORAI1 Proteins in Intact Cells Via Correlative Light and Electron Microscopy. Microsc. Microanal. 22:902–912.

14. Alansary, D., D. B. Peckys, B. A. Niemeyer, and N. de Jonge. 2020. Detecting single ORAI1 proteins within the plasma membrane reveals higher order channel complexes. J. Cell Sci. 133: 240358-1-12.

15. Dahmke, I. N., A. Verch, J. Hermannsdorfer, D. B. Peckys, R. S. Weatherup, S. Hofmann, and N. de Jonge. 2017. Graphene Liquid Enclosure for Single-Molecule Analysis of Membrane Proteins in Whole Cells Using Electron Microscopy. ACS Nano 11:11108–11117.

16. Zhou, Y., R. M. Nwokonko, X. Cai, N. A. Loktionova, R. Abdulqadir, P. Xin, B. A. Niemeyer, Y. Wang, M. Trebak, and D. L. Gill. 2018. Cross-linking of Orai1 channels by STIM proteins. Proc. Natl. Acad. Sci. 115:E3398–e3407.

17. Gwack, Y., S. Srikanth, S. Feske, F. Cruz-Guilloty, M. Oh-hora, D. S. Neems, P. G. Hogan, and A. Rao. 2007. Biochemical and functional characterization of Orai proteins. J. Biol. Chem. 282:16232–16243.

18. Kim, J. H., S.-R. Lee, L.-H. Li, H.-J. Park, J.-H. Park, K. Y. Lee, M.-K. Kim, B. A. Shin, and S.-Y. Choi. 2011. High Cleavage Efficiency of a 2A Peptide Derived from Porcine Teschovirus-1 in Human Cell Lines, Zebrafish and Mice. PLOS ONE 6:e18556.

19. Peckys, D. B., and N. de Jonge. 2015. Studying the stoichiometry of epidermal growth factor receptor in intact cells using correlative microscopy. J. Vis. Exp. 103:e53186.

20. Gilkey, J. C., and L. A. Staehelin. 1986. Advances in ultrarapid freezing for the preservation of cellular ultrastructure. J. Electron Microsc. Tech. 3:177–210.

21. Stanly, T. A., M. Fritzsche, S. Banerji, E. García, J. B. de la Serna, D. G. Jackson, and C. Eggeling. 2016. Critical importance of appropriate fixation conditions for faithful imaging of receptor microclusters. Biol. Open 5:1343–1350.

22. Kusumi, A., and K. Suzuki. 2005. Toward understanding the dynamics of membraneraft-based molecular interactions. Biochim. Biophys. Acta 1746:234–251.

23. Tanaka, K. A. K., K. G. N. Suzuki, Y. M. Shirai, S. T. Shibutani, M. S. H. Miyahara, H. Tsuboi, M. Yahara, A. Yoshimura, S. Mayor, T. K. Fujiwara, and A. Kusumi. 2010. Membrane molecules mobile even after chemical fixation. Nat. Methods 7:865–866.

24. Textor, M., and N. de Jonge. 2018. Strategies for Preparing Graphene Liquid Cells for Transmission Electron Microscopy. Nano Lett. 8: 3313–3321.

25. Blach, P., S. Keskin, and N. de Jonge. 2020. Graphene Enclosure of Chemically Fixed Mammalian Cells for Liquid-Phase Electron Microscopy. J. Vis. Exp.

26. Peckys, D. B., U. Korf, and N. de Jonge. 2015. Local variations of HER2 dimerization in breast cancer cells discovered by correlative fluorescence and liquid electron microscopy. Sci. Adv. 1:e1500165.

27. Peckys, D. B., C. Quint, and N. Jonge. 2020. Determining the Efficiency of Single Molecule Quantum Dot Labeling of HER2 in Breast Cancer Cells. Nano Lett. 20:7948–7955.

28. Peckys, D. B., and N. de Jonge. 2014. Liquid scanning transmission electron microscopy: imaging protein complexes in their native environment in whole eukaryotic cells. Microsc. Microanal. 20:346–365.

29. Scrimgeour, N., T. Litjens, L. Ma, G. J. Barritt, and G. Y. Rychkov. 2009. Properties of Orai1 mediated store-operated current depend on the expression levels of STIM1 and Orai1 proteins. J. Physiol. 587:2903–2918.

30. Hoover, P. J., and R. S. Lewis. 2011. Stoichiometric requirements for trapping and gating of Ca2+ release-activated Ca2+ (CRAC) channels by stromal interaction molecule 1 (STIM1). Proc. Natl. Acad. Sci. 108:13299–13304.

31. Brandman, O., J. Liou, W. S. Park, and T. Meyer. 2007. STIM2 Is a Feedback Regulator that Stabilizes Basal Cytosolic and Endoplasmic Reticulum Ca2+ Levels. Cell 131:1327–1339.

32. Peckys, D. B., C. Stoerger, L. Latta, U. Wissenbach, V. Flockerzi, and N. de Jonge. 2017. The stoichiometry of the TMEM16A ion channel determined in intact plasma membranes of COS-7 cells using liquid-phase electron microscopy. J. Struct. Biol. 199:102–113.

33. Hsieh, T.-S., Y.-J. Chen, C.-L. Chang, W.-R. Lee, and J. Liou. 2017. Cortical actin contributes to spatial organization of ER–PM junctions. Mol. Biol. Cell 28:3171–3180.

34. Orci, L., M. Ravazzola, M. Le Coadic, W.-w. Shen, N. Demaurex, and P. Cosson. 2009. STIM1-induced precortical and cortical subdomains of the endoplasmic reticulum. Proc. Natl. Acad. Sci. 106:19358.

35. Peckys, D. B., U. Korf, S. Wiemann, and N. De Jonge. 2017. Liquid-phase electron microscopy of molecular drug response in breast cancer cells reveals irresponsive cell subpopulations related to lack of HER2 homodimers. Mol. Biol.Cell 28:3193–3202.

36. Peckys, D. B., D. Hirsch, T. Gaiser, and N. de Jonge. 2019. Visualisation of HER2 homodimers in single cells from HER2 overexpressing primary formalin fixed paraffin embedded tumour tissue. Mol. Med. 25:42.

37. Perni, S., J. L. Dynes, A. V. Yeromin, M. D. Cahalan, and C. Franzini-Armstrong. 2015. Nanoscale patterning of STIM1 and Orai1 during store-operated Ca2+ entry. Proc. Natl. Acad. Sci. 112:E5533–E5542.

38. Kusumi, A., and K. Suzuki. 2005. Toward understanding the dynamics of membraneraft-based molecular interactions. Biochim. Biophys. Acta 1746:234–251.

39. Huebinger, J., J. Spindler, K. J. Holl, and B. Koos. 2018. Quantification of protein mobility and associated reshuffling of cytoplasm during chemical fixation. Sci. Rep. 8:17756.

40. Demuro, A., A. Penna, O. Safrina, A. V. Yeromin, A. Amcheslavsky, M. D. Cahalan, and I. Parker. 2011. Subunit stoichiometry of human Orai1 and Orai3 channels in closed and open states. Proc. Natl. Acad. Sci. 108:17832–17837.

41. Katz, Z. B., C. Zhang, A. Quintana, B. F. Lillemeier, and P. G. Hogan. 2019. Septins organize endoplasmic reticulum-plasma membrane junctions for STIM1-ORAI1 calcium signalling. Sci. Rep. 9:10839.

42. Hartzell, C. A., K. I. Jankowska, J. K. Burkhardt, and R. S. Lewis. 2016. Calcium influx through CRAC channels controls actin organization and dynamics at the immune synapse. eLife 5:e14850.

43. Yadav, R., and H. P. Lu. 2018. Revealing dynamically-organized receptor ion channel clusters in live cells by a correlated electric recording and super-resolution singlemolecule imaging approach. Phys. Chem. Chem. Phys. 20:8088–8098.

44. Tobin, S. J., D. L. Wakefield, L. Terenius, V. Vukojevic, and T. Jovanovic-Talisman. 2019. Ethanol and Naltrexone Have Distinct Effects on the Lateral Nano-organization of Mu and Kappa Opioid Receptors in the Plasma Membrane. ACS Chem. Neurosci. 10:667–676.

45. Needham, S. R., M. Hirsch, D. J. Rolfe, D. T. Clarke, L. C. Zanetti-Domingues, R. Wareham, and M. L. Martin-Fernandez. 2013. Measuring EGFR Separations on Cells with ~10 nm Resolution via Fluorophore Localization Imaging with Photobleaching. PLoS ONE 8:e62331.

46. Parker, K., P. Trampert, V. Tinnemann, D. B. Peckys, T. Dahmen, and N. de Jonge. 2018. Linear chains of HER2 receptors found in the plasma membrane using liquid-phase electron microscopy. Biophys. J. 115:503–513.

47. Hagner, K., S. Setayeshgar, and M. Lynch. 2018. Stochastic protein multimerization, activity, and fitness. Phys. Rev. E 98:062401.

48. Hashimoto, K., and A. R. Panchenko. 2010. Mechanisms of protein oligomerization, the critical role of insertions and deletions in maintaining different oligomeric states. Proc. Natl. Acad. Sci. 107:20352–20357.

49. Lopez-Guerrero, A. M., P. Tomas-Martin, C. Pascual-Caro, T. Macartney, A. Rojas-Fernandez, G. Ball, D. R. Alessi, E. Pozo-Guisado, and F. J. Martin-Romero. 2017. Regulation of membrane ruffling by polarized STIM1 and ORAI1 in cortactin-rich domains. Sci. Rep. 7:383.

50. Leverrier-Penna, S., O. Destaing, and A. Penna. 2020. Insights and perspectives on calcium channel functions in the cockpit of cancerous space invaders. Cell Calcium:102251.

51. Lewis, R. S. 2020. Store-Operated Calcium Channels: From Function to Structure and Back Again. Cold Spring Harb. Perspect. Biol. 12:a035055.

52. Shaw, P. J., B. Qu, M. Hoth, and S. Feske. 2013. Molecular regulation of CRAC channels and their role in lymphocyte function. Cell. Mol. Life Sci. 70:2637–2656.

