## Supplementary Figures for "Supra-molecular assemblies of ORAI1 at rest precede local accumulation into punctae after activation"

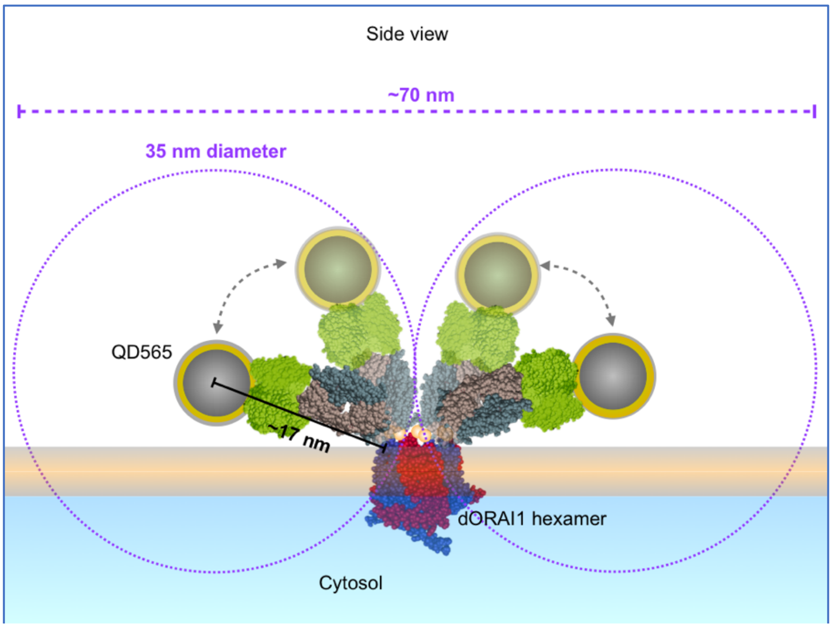


*Figure S1*. Scheme illustrating the most distant positioning of two QD labels (shown here are the smaller QD565) bound to the same hexameric ORAI1 channel. The max. label-to-protein distance is ~17 nm, thus similar as found for the QD655 (shown in Fig.1a). Our image processing tool enlarges all detected labels to circles with a diameter of 35 nm (as shown by the two dotted purple circles), each indicating the possible max. range for the location of the underlying, labeled ORAI1 protein. In the depicted, most extreme position, both circles would still touch each other, and thus be detected as a cluster. For labels bound to the same ORAI1 channel, the major axis of a label cluster cannot exceed 70 nm, i.e. twice the diameter of both label circles. It follows that all detected cluster >70 nm must indicate more than one underlying ORAI1 channel. Note, that all involved particles and proteins are drawn to scale, as in Fig. 1.

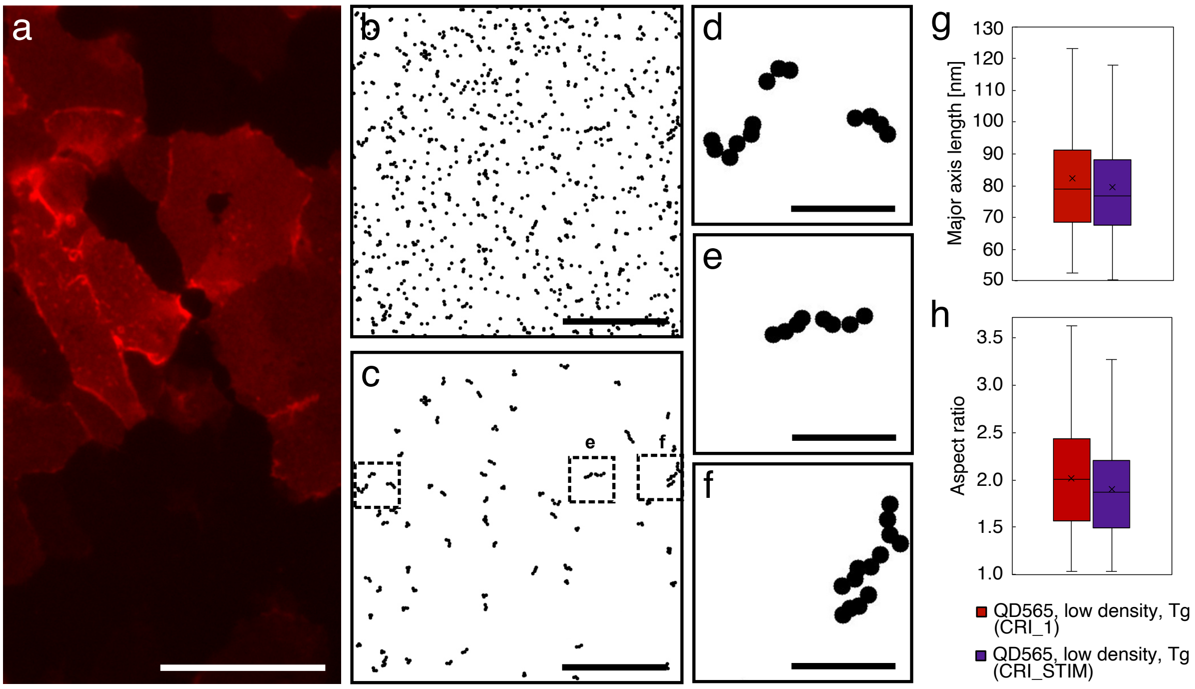


*Figure S2.* ORAI1 distributions in HEK cells lacking STIM1 and 2 proteins (CRI_STIM cell line) proving that linear, supramolecular ORAI1 arrangements do not depend on the presence of STIM proteins. (a) Fluorescence image of cells expressing ORAI1 after 15 min of Tg incubation, fixation and labeling with QD655 show a similar homogenous ORAI1 distribution throughout the plasma membrane as found in resting HEK cells lacking ORAI1 and 2, but expressing endogenous STIM (CRI_1) (compare to Fig. 1c). (b) The processed version of an exemplary scanning transmission electron microscopy (STEM) image, showing all label positions as 35 nm circles, reveals a scattered distribution of single and clustered labels. (c) The same image after cluster analysis, showing only clusters >2, with similar dimensions and linear arrangements as found for cluster >2 in resting cells (compare to Fig. 2b and d). The boxed regions are shown enlarged in (d-f), disclosing linear arrangements of ORAI1. (g, h) Quantitative analysis of maximal dimension (g) and aspect ratio (h) of cluster >2 in low expressing CRI_1 cells after maximal SOCE-activation with Tg (red, 277 cluster) and in low expressing CRI_STIM cells after Tg incubation (purple, 431 cluster). The maximal dimension differed marginally by3% ($p<0.054$), aspect ratio (AR) was slightly higher (7%) in the presence of STIM proteins ($p<0.0111$). Scale bar in a = 50 μm, in b and c = 1 μm, and in d – f = 200 nm.

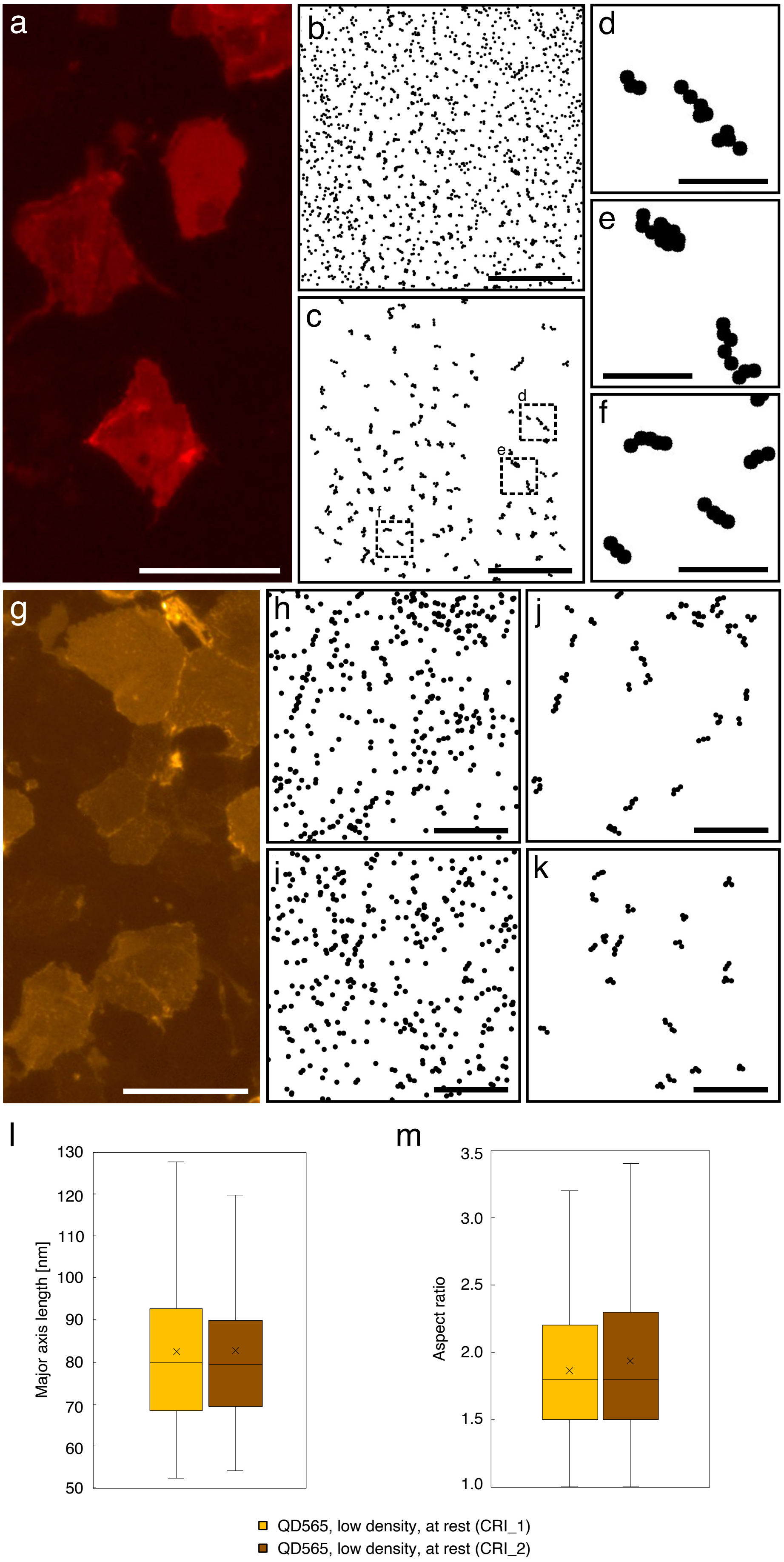
a
b
c
d
e
f
g
h
i
j
k
l
m
130
120
110
100
90
80
70
60
50
Major axis length [nm]
3.5
3.0
2.5
2.0
1.5
1.0
Aspect ratio
QD565, low density, at rest (CRI_1)
QD565, low density, at rest (CRI_2)

*Figure S3.* Control experiments were performed with a HEK cell line lacking endogenous ORAI1-3 proteins (CRI_2) to test a possible influence of ORAI3. These experiments were also performed with smaller QD565 labels to study a possible effect of the QD label size on the spatial dimensions and shapes of the detected labeled ORAI1 clusters. (a) Fluorescence image of CRI_2 cells at rest with QD655-labeled ORAI1 show a homogenous staining of the plasma membrane. (b) The processed version of an exemplary STEM image, showing all label positions as 35 nm circles, reveals a typical scattered distribution of single and clustered labels. (c) The same image after cluster analysis, showing only clusters >2, with similar dimensions and linear arrangements as found for cluster >2 in resting cells (compare to Fig. 2b and d). The boxed regions showing typical linear arrangements of ORAI1 are shown enlarged in (d-f). (g) Fluorescence image of CRI_2 cells at rest labeled with QD565, shows also homogenous plasma membrane staining. (h, i) Two typical processed STEM images showing all detected QD565 labels as 35 nm circles, demonstrating similarly scattered distributions of labels in monodispersed or clustered conformations as found in the CRI_1 cell line, and with the larger QD655 labels. Note that the magnification used to record the STEM images of QD565 labeled cells was usually set 2-fold higher, as for the QD655 labels, as needed to resolve their smaller size. (j, k) Corresponding images after cluster analysis showing only cluster >2, reveal similar linear ORAI1 cluster >2 as found in CRI_1 and CRI_2 cells labeled with QD655. (l, m) Quantitative analysis of max. dimension (l) and AR (m) of QD565-labeled ORAI1 cluster >2 in resting, low expressing CRI_1 (249 cluster*)*, and CRI_2 cells (209 cluster). No significant difference was found between the two cell lines ($p$ = 0.85 for the length, $p$ = 0.14 for AR). Scale bar in a = 50 µm, in b and c = 1 µm, and in d – f = 200 nm.

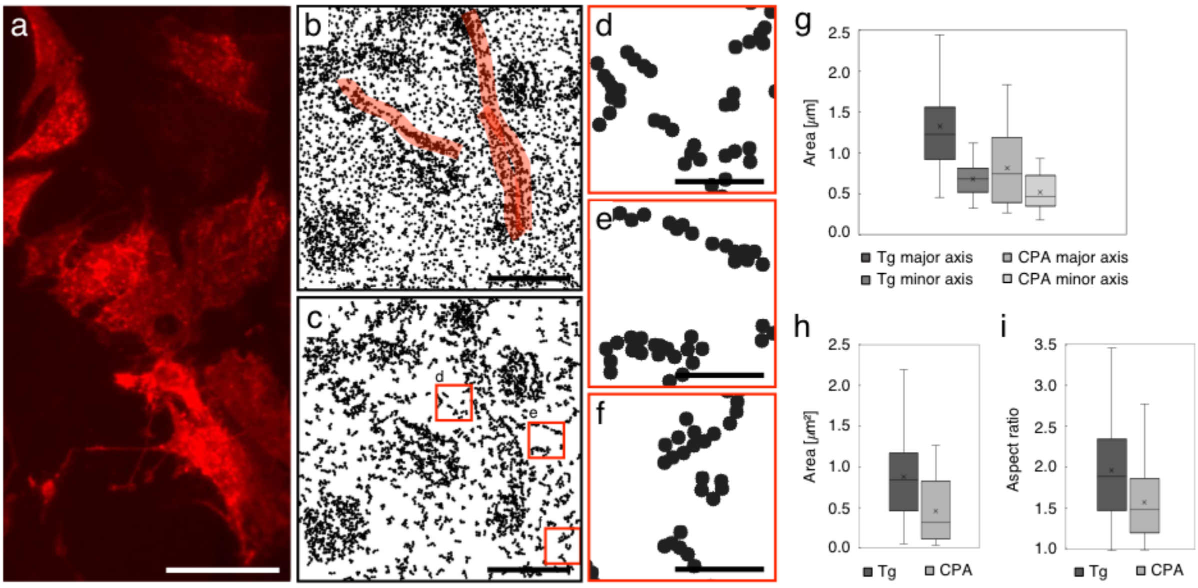


*Figure S4*. Control experiments were performed to test possible differences of CPA-induced, compared to Tg-induced SOCE-activation, both performed in CRI_1 cells co-expressing STIM1 with ORAI1 in a 3:1 ratio. (a) Fluorescence image of QD655-labeled ORAI1 shows a dotted staining of the plasma membrane, similar to the situation found in Tg-activated cells (see Fig. 3a). (b, c) Exemplary STEM image from a high expressing cell after CPA-induced SOCE-activation, processed to show all label positions as 35 nm circles (b), respect. in (c) only those labels found in clusters >2. Several roundish ORAI1 accumulation areas, as well as long strands (high-lightened with transparent red lines) are seen. In (c) three regions outside of punctae are marked, the cluster found in these regions are shown in detail in (d-f). Their elongated and linear shapes, as well as their length are similar to cluster >2 found in cells at rest and in cells activated with Tg, outside of the punctae. (g) Comparison of the major and minor axis dimensions of punctae after Tg (dark gray, 44 punctae) and after CPA activation (light grey, 36 punctae), shows that CPA lead to smaller punctae. (h, i) Quantitative analysis of the area size distribution (h) and the AR (i) of punctae in the Tg and CPA activation reveals that CPA-induced punctae are approx. 2-times smaller and significantly rounder than punctae found after Tg activation. All differences between CPA- and Tg-induced punctae parameter are highly significant ($p < 0.001$). Scale bar in a = 50 µm, in b and c = 1 µm, and in d – f = 200 nm.

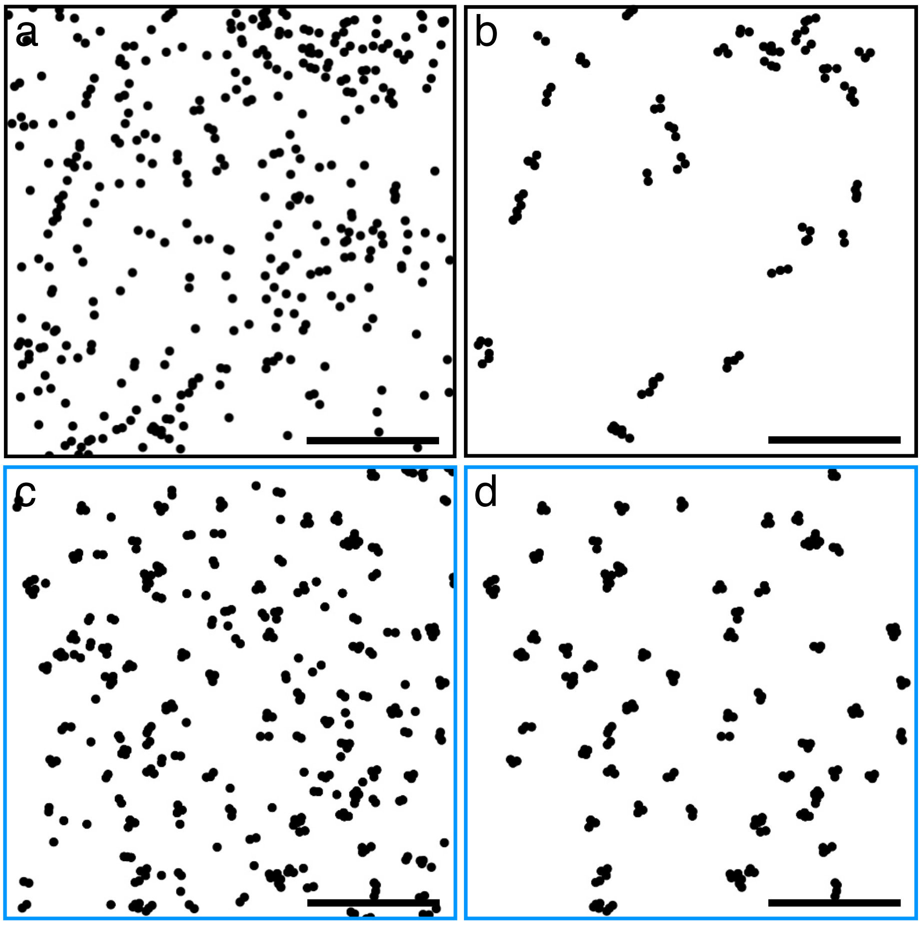


*Figure S5.* Example of a processed STEM image of ORAI1 labeled with QD565 on a low expressing cell at rest (a), showing all detected QD labels, and the corresponding simulated label distribution (c), based on the same image size and number of detected labels, bound with a 30% labeling efficiency to randomly distributed, hexameric ORAI1. The results of the cluster analysis are shown for the STEM data in (b), revealing mostly elongated and linear cluster >2, and for the simulation in (d), showing mostly smaller and rounder cluster >2. Scale bars = 500 nm.

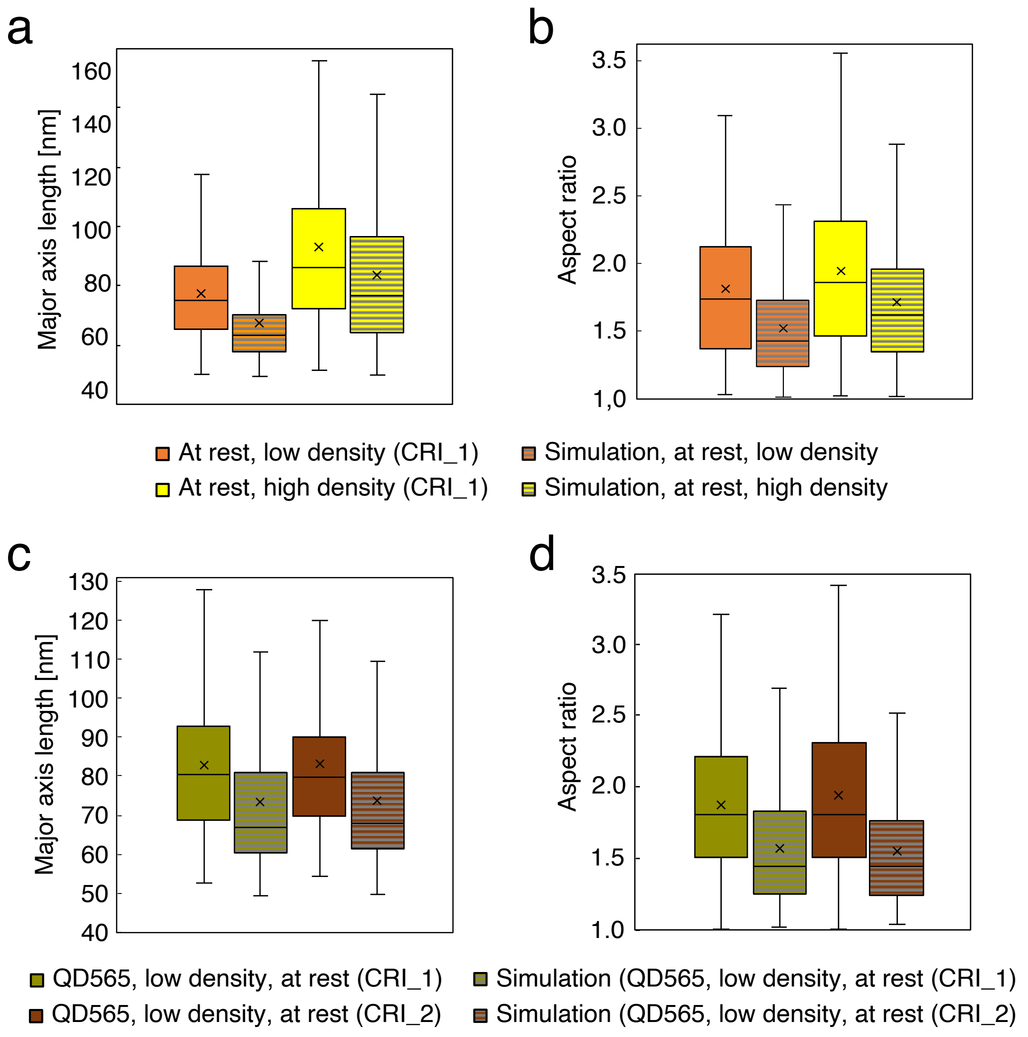


*Figure S6*. Quantitative results of the cluster analysis comparing between the STEM image data sets of QD655 and QD565 labeled ORAI1 and the corresponding simulations of the label density-matching distributions, for resting cells of the CRI_1 and CRI_2 cell line. (a) Cluster >2 detected in the data from low expressing cells at rest (orange) are larger than those in their corresponding simulations (orange/grey striped). 75% of clusters found in the STEM images from cells exceed the 70 nm limit (the max. dimension for cluster of labels bound to the same hexameric ORAI1 channel), whereas 75% of clusters found in the simulations are smaller than 70 nm. High expressing cells (yellow) have larger dimensions than low expressing cells, due to a higher share in coalescing clusters, being measured as one single cluster. Also, for the high expressing cells, comparison with the corresponding simulation (yellow/grey striped) shows that the cluster dimensions in STEM images are significantly larger. (b) The results from measurements of the cluster AR show similar results with higher values for cluster >2 in STEM images than those found in the matching simulations. (c and d) Quantitative cluster analysis for QD565 labeled cells of the main cell line (CRI_1, green, 249 cluster) and the control cell line lacking ORAI1-3 (CRI_2, brown, 208 cluster), and their corresponding simulations (green/grey striped for CRI-1, respect. brown/grey striped for CRI_2), confirm the results of the QD655 data, namely higher values for cluster max. dimensions and AR in STEM images, compared with density-matching simulations. For all comparisons between image data and their respect. simulations $p < 0.001$.

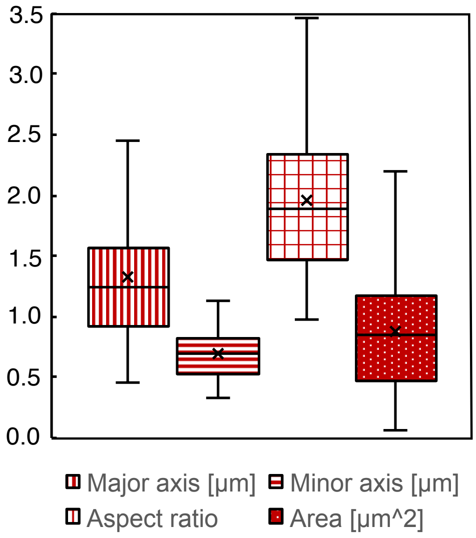


*Figure S7*. Quantitative measurements of punctae dimensions, shape and area size in max. SOCE-activated cells (with 3:1 co-expression of STIM1/ORAI1) after 15 min of Tg. The analysis included all visually identifiable ORAI1 accumulation areas (n = 44 punctae), being completely depicted in STEM images of QD655-labeled CRI-1 cells (N = 11).

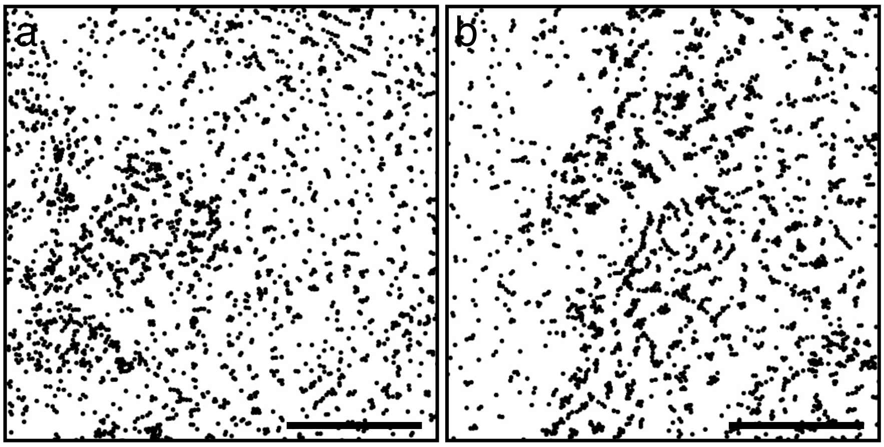


*Figure S8*. Two examples of processed STEM data (all labels are depicted as black circles) from QD655 labeled cells after submax. SOCE activation (1:1 co-expression of STIM1/ORAI1, after 15 min of Tg). Accumulation of ORAI1 clusters in punctae can be discerned, but compared to images recorded after max. activation, the ORAI1 density within the accumulation areas is lower (compare to Fig. 3b). The lower density enables a better discrimination of 2-D structures. ORAI1 seemed to have accumulated by formation of larger clusters, often consisting of two aligned elongated ORAI1 cluster >2, leaving more free space between these larger cluster than seen in max. activated cells. The borders of these punctae regions appear blurrier than after max. activation. Scale bars = 1 µm.
